# AICAr inhibition of cardiomyocyte autophagy promotes p62-dependent NRF2 expression and protection against doxorubicin toxicity

**DOI:** 10.1101/2025.08.20.671370

**Authors:** Erin K. Fassett, Bernd Mayer, John T. Fassett

## Abstract

Doxorubicin is an effective cancer chemotherapeutic, but its use is complicated by cardiotoxic side-effects. 5-amino-4-imidazolecarboxamide ribonucleoside (AICAr) is a widely used pharmacological activator of adenosine monophosphate activated kinase (AMPK), but also exerts AMPK-independent actions that may have unrealized therapeutic potential. Here, we identified a novel mechanism by which pretreatment with AICAr protects neonatal rat cardiomyocytes against doxorubicin toxicity. Despite increasing AMPK^Thr172^ and ULK1^Ser555^ phosphorylation, AICAr suppressed cardiomyocyte LC3 lipidation and caused accumulation of the autophagy receptor, p62 SQST1, through an adenosine kinase (ADK)-dependent, AMPK-independent mechanism. The accumulation of p62 was associated with increased expression and transcriptional activity of NRF2, as well as decreased doxorubicin-induced reactive oxygen species and cell death. Notably, AICAr-induced increase of NRF2, antioxidant gene expression, and doxorubicin resistance were blocked by RNAi depletion of p62, indicating that the protective effects of AICAr rely upon the secondary effects of autophagy inhibition, rather than autophagy inhibition *per se*. While doxorubicin treatment alone did not affect cardiomyocyte LC3 lipidation, it did significantly decrease p62 levels and diminish NRF2 nuclear localization. Pretreatment with AICAr to provide surplus p62 and nuclear NRF2 diminished the impact of these doxorubicin effects. Importantly, MCF7 breast cancer cells, which poorly express ADK, were not protected by AICAr pretreatment and instead were sensitized to doxorubicin-induced cell death. These findings raise the possibility that differences in ADK expression between cardiomyocytes and breast cancer cells might be exploited by pretreatment with AICAr or similar ADK-dependent drugs to provide dual benefits in doxorubicin therapy.

## Introduction

Doxorubicin (Dox), an anthracycline antibiotic, is a cytotoxic chemotherapeutic agent effective against many cancers, including leukemia, lymphoma, ovary, bladder, thyroid and breast cancer [18]. Doxorubicin acts on cancer cells by intercalating between DNA base pairs and inhibiting topoisomerase2α, resulting in single and double-stranded breaks, initiation of apoptosis, and inhibition of DNA and RNA synthesis [10]. Doxorubicin also forms reactive oxygen species (ROS) which results in cytotoxicity to cancer cells secondary to lipid peroxidation of cell membranes [21]. Unfortunately, dose-dependent acute and chronic cardiotoxic effects limit its use as a chemotherapeutic. Despite decades of research, the underlying molecular mechanisms of doxorubicin cardiotoxicity have yet to be completely understood [13]. While oxidative stress has been implicated as a primary cause of dox-induced cardiotoxicity, there are no clinical studies in humans showing that exogenous antioxidant treatment is effective against Doxorubicin cardiotoxicity [13, 42]. Additional suspected contributors to doxorubicin cardiotoxicity are ferroptosis, inhibition of Topoisomerase2ß, disruption of cardiac metabolic homeostasis via decreased AMPK activity, and perturbations of autophagy [29]. Currently only one drug, dexrazoxane, an iron chelator, has been approved for clinical use by the FDA for treatment of doxorubicin toxicity. However, dexrazoxane’s use is limited by serious adverse reactions such as myelotoxicity [29]. Thus, the identification of new approaches to protect against doxorubicin-induced cardiotoxicity without diminishing chemotherapeutic efficacy remains clinically relevant.

Activation of endogenous cardioprotective signaling pathways prior to doxorubicin treatment may help shield against doxorubicin-induced cell damage. One cardioprotective response that has shown promise in protection against doxorubicin cardiotoxicity is the KEAP1/ NRF2/ ARE (Kelch-Like ECH Associated Protein 1/Nuclear factor erythroid 2-related factor 2/Antioxidant Response Elements) signaling pathway [52]. During basal conditions, KEAP1 limits NRF2 expression by promoting its ubiquitination and proteasomal degradation [3]. Disruption of the KEAP-1-NRF2 interaction under oxidative stress conditions increases NRF2 levels [51], which induces transcription of detoxification enzymes and antioxidants such as catalase (CAT), NADPH Quinone Dehydrogenase 1 (NQO1), and heme oxygenase-1 (HO-1). Importantly, exogenous NRF2 expression [26] or drugs that elevate NRF2 activity by disrupting KEAP1-NRF2 interaction can attenuate cardiac damage by doxorubicin [28, 52].

The autophagy receptor, p62 SQSTM-1 (sequestosome 1), plays a key role in NRF2 signaling by binding to KEAP1 and interfering with NRF2 ubiquitination and degradation [3, 23, 24]. Additionally, NRF2 promotes p62 transcription, so that increased NRF2 activity creates a positive feedback loop by enhancing p62-dependent inhibition of KEAP1 [16]. Thus, transient inhibition of autophagy to elevate p62 levels could set off a sustained increase in NRF2 signaling [17] that protects against oxidative stress and other cell damaging effects of doxorubicin.

Activation of the metabolic sensor, adenosine monophosphate-activated protein kinase (AMPK) has also been shown to attenuate doxorubicin cardiotoxicity [27, 47]. However, a recent clinical trial demonstrated that the AMPK-activating drug, metformin, did not significantly diminish doxorubicin-induced cardiac injury in breast cancer patients [35], suggesting activation of AMPK alone may be insufficient for cardioprotection.

Additionally, drugs that enhance AMPK activity alone may promote autophagic degradation of p62, thereby diminishing p62-mediated preservation of NRF2 signaling. 5-amino-4-imidazolecarboxamide ribonucleoside (AICAr) is an endogenous purine synthesis intermediate that has found widespread use as a pharmacological agent for investigating the role(s) of AMPK *in vitro* and *in vivo.* AICAr enters the cells through nucleoside transporters and becomes phosphorylated by adenosine kinase (ADK) into 5-amino-4-imidazolecarboxamide ribonucleotide (AICAR), also known as ZMP. AICAr effects are therefore enhanced in cells expressing high levels of ADK. While ZMP is approximately 50 times less potent than 5’adenosine-monophosphate (AMP) in AMPK activation, it is slowly metabolized, so that it accumulates to high concentrations in the cytoplasm (up to 10mM) and strongly activates AMPK [7]. Besides activating AMPK, however, AICAr has numerous AMPK-independent effects which may have unrealized therapeutic potential [50]. One such AICAr effect is inhibition of autophagy. Despite increasing AMPK activity, AICAr has been shown to suppress autophagy in a variety of cell types [15, 40, 49]. Because autophagy inhibition causes p62 accumulation and subsequently increases NRF2 [23], AICAr has potential to simultaneously elicit protective effects of both AMPK and p62/NRF2 signaling. The effects of AICAr on cardiomyocyte autophagy however, and how this might impact doxorubicin-induced cardiotoxicity, have not been described.

Here, we investigated the effects of AICAr on autophagy, p62/NRF2 signaling and doxorubicin tolerance in cardiomyocytes and MCF7 breast cancer cells. Our findings identify a novel ADK-dependent mechanism by which AICAr pretreatment transiently inhibits autophagy to induce p62-dependent protection of cardiomyocytes against doxorubicin toxicity.

## Materials and Methods

### Animal Care and Tissue Collection

Animal care complied with the Austrian law on experimentation with laboratory animals (last amendment, 2013) based on the European Union guidelines for the Care and the Use of Laboratory Animals (European Union Directive 2010/63/EU) and is reported in accordance with the ARRIVE guidelines for reporting experiments involving animals [20, 32].

### Neonatal Cardiomyocyte Culture and Treatments

Neonatal cardiomyocytes (NRVMs) were isolated from 3–4-day old neonatal rats as previously described [8] and plated at 8×10^4^ cells/cm^2^ on 0.1% gelatin-coated plates in low glucose(1g/L) DMEM containing 10% FCS. After attachment overnight, 0.1mM bromodeoxyuridine was added for an additional 24 hours to prevent fibroblast proliferation. Where indicated, cells were treated with RNAi p62 RNAi (Dharmacon; cat#L-089365-02-0005), or non-targeting control RNAi (Dharmacon; cat#D-001810-10-20) as previously described [9]. For adenoviral expression of AMPKα2, NRVMs were incubated overnight with the indicated MOI of adenovirus. Experiments were performed 48 hours later with fresh medium. NRVMs were treated with AICAr (MCE cat#HY-13417/CS-1951), ABT-702 (MCE cat#HY-103161), Bafilomycin (Santa Cruz cat#SC-201550A), and/or Doxorubicin hydrochloride (Tocris; cat#2252), at the indicated dosages and time periods.

### MCF7 Cells Culture and Treatment

MCF7 cells, kindly donated by the Groschner lab, Medical University of Graz, were grown in DMEM with (3g/L) glucose, (55mg/L) pyruvate supplemented with insulin-transferrin-selenium (ITS) (Gibco; cat#41400-045), Penicillin-Streptomycin, and 10% FBS. Cells were maintained in a humidified atmosphere of 5% CO2 at 37 °C. Cell media was changed every 2-3 days unless otherwise noted. Once MCF7 cells reached ∼70% confluency, they were treated with AICAr, Doxorubicin, and/or Bafilomycin in fresh media at the indicated dosages and times.

### Western Blot Analysis of Cell Lysates

Cell lysates were collected in 1x SDS loading buffer containing protease inhibitors (Boehringer-Mannheim; cat# 04 693 132 001) and phosphatase inhibitors (Boehringer-Mannheim; cat# 04 906 837 001), heated to 95°C for 1 minute and frozen at −80°C until analysis. Cell lysates were reduced with 0.1M DTT for 30 minutes at room temperature before analysis by Western blotting. Equal amounts of protein were separated by SDS/PAGE and then transferred to 0.2 µm PVDF or nitrocellulose membranes (Amersham).

Membranes were blocked in 5%NF milk for 1 hour before adding primary antibodies overnight at 4°C. Antibodies for LC3A/B (cat# #12741), Phospho-AMPK^Thr172^ (cat#2535) and Phospho-ULK ^Ser555^ (cat#5869T) were from Cell Signaling Technology. Antibodies for p62 (cat#18420-1-AP), NRF2 (cat#16396-1-AP), GAPDH (cat#60004-1-Ig), Sarcomeric actin (cat#66125-1-Ig), and ßactin (cat#66009-1-Ig) were from Proteintech. Membranes were incubated with secondary antibodies to rabbit (cat#7074S) or mouse (cat#7076S) from Cell Signaling Technology for 1 hour at room temperature. Chemiluminescent signals were detected using Quantum (cat#541015) or Western Bright (cat#541005) from Advansta and recorded using Fusion imaging system (Vilber Lourmat, Germany). Quantification was performed using ImageJ. Band intensities were normalized to constitutively expressed loading controls (sarcomeric actin, β-actin or GAPDH) on the same blot (with the exception of ADK expression in figure 8, where expression of GAPDH in the different cell/tissue types widely differed). Autophagic flux was determined by measuring the difference between LC3-II levels in bafilomycin and non-bafilomycin treated cells in the same experiment.

### HPLC analysis of ZMP

NRVMs in 12-well plates were treated with vehicle, AICAr, or AICAR+ABT for 3 hours and washed with PBS before adding 100 µl 0.4 M perchloric acid (PCA). After 5 min on ice, the supernatant of the samples was removed and neutralized with 75 µl 2.5 M KOH in 1 M K2HPO4, allowed to stand for 5 min at 4 °C, and centrifuged at 20 000 g for 10 min at 4 °C. Supernatant was removed for analysis. ZMP was quantified by HPLC (Agilent 1260 Infinity with diode array detection) on a reversed phase Poroshell Bonus RP column (Agilent Technologies). Aliquots of 6 µl sample were injected onto the column and analyzed at a flow rate of 1.0 ml/min and a column temperature of 35 °C using a mobile phase gradient of 0-8 minutes buffer A (90 mM KH2PO4, 10mM K2HPO4, 10 mM TBAHS, pH 6.0), 8-14 minutes 96% buffer A 4% methanol, and 14-20 minutes 88% buffer A 12% methanol. ZMP was calibrated in the range of 1-100 µM based on the area of the peak detected at retention time 1.57 min and analyzed with Open LAB software. Protein concentration from the cells was measured using BCA assay after dissolving in 5% SDS, 0.1N NaOH. Values are expressed as nmoles/mg protein.

### RNA Isolation-RT-qPCR

Total RNA was isolated from NRVMs following treatment plus or minus CTR RNAi, p62 RNAi, and AICAr using TRIzol Reagent (Invitrogen cat#15596026). RNA yield and purity were determined using NanoDrop Spectrophotometer. cDNA was synthesized from total RNA using the High Capacity cDNA Reverse Transcription Kit (Applied Biosystems). Quantitative PCR was performed with SsoAdvanced Universal SYBR^®^ Green Supermix (BIO-RAD, cat. # 172-5271). Primers for *18s*, *p62, NRF2*, *NQO1*, *HO1*, and *SOD1* were from Invitrogen (**Table 1**). Data was acquired from 3 independent experiments with 2-3 biological replicates per condition.

**Table 1:**
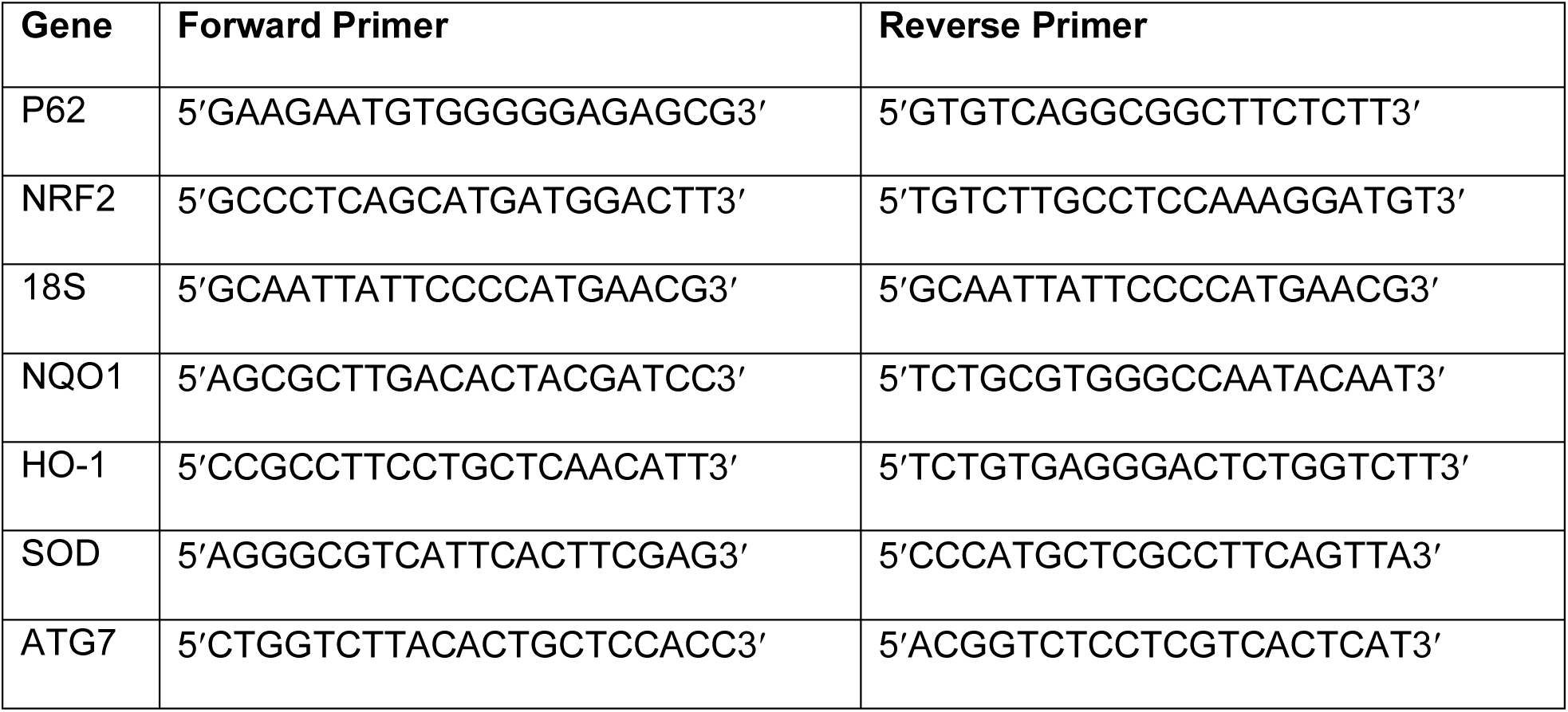
Sequence of Primers.

### XTT Assay

Cell viability was assessed with Cayman’s XTT cell proliferation assay (cat#10010200). In brief, cells were seeded on a 96-well plate in 100ul of media plus FCS, maintained in 5% CO2 at 37°C and treated as described in the text. After 6 hours of doxorubicin treatment, media was replaced, and cells were incubated for an additional 24 hours. Cells were then treated with 10ul of prepared XTT Mixture and incubated for an additional 2 hours. The absorbance was measured on a microplate reader at a wavelength of 450 nm. Cell viability measurements are normalized to untreated controls, which are set at 1. For comparing the effects of AICAr on doxorubicin toxicity in control RNAi and p62 RNAi-treated cells, percent cell death was used, since p62 RNAi treatment alone decreased cell growth and XTT values. Box and Whisker plots represent data from at least 3 independent experiments each containing 3-6 biological replicates per condition.

### Luciferase Assay

NRVMs in 48 well plates were co-transfected overnight with an NRF2 luciferase reporter plasmid, pGL4.27_ ARE/NRF2-SPE [14] and a constitutive renilla-luciferase reporter, pLX313-Renilla [12] at a 10:1 ratio using Turbofect (ThermoFisher, cat# RO534) according to the manufacturer’s instructions. One day after transfection, cells were treated with 500µM AICAr for 24 hours. Firefly and renilla luciferase activities were measured on a fluorescent plate reader (Promega, Glo-Max) using the Dual-Glow Luciferase Assay System (Promega, cat# E2920) according to the manufacturer’s instructions. pGL4.27_ARE/NRF2-SPE was a gift from Michael Ristow (Addgene plasmid # 177775; http://n2t.net/addgene:177775; RRID:Addgene_177775). pLX313-renilla luciferase was a gift from William Hahn & David Root (Addgene plasmid # 118016; http://n2t.net/addgene:118016; RRID:Addgene_118016).

### Detection of ROS via DCFDA Assay

Carboxy-H_2_DCFDA is non-fluorescent until it is oxidized by the presence of ROS, at which point it is detectable as green fluorescence. NRVMs were seeded on an ibidi µ-plate (#80826) in 200µl media plus FCS. Cells were incubated in the presence or absence of 500µM AICAr for 24 hours, after which media was replaced with 200µl serum-free media. Next, cells were treated with Carboxy-H_2_DCFDA (MCE cat# HY-D0940/CS-7539) at a final concentration of 5µM in 200µl serum-free media and incubated at 37°C in 5% CO2 for 30 minutes. Cells were then washed 2x with HBSS and 200µl media containing doxorubicin or vehicle was added for 5 hours. Green fluorescence was imaged using the GFP fluorescence filter and quantitated using ImageJ.

### Immunofluorescence

Cells were plated on glass coverslips coated with 0.1% gelatin or μ-Slide 8-well plates from Ibidi (cat#80826). After the indicated treatments and time periods, cells were fixed in methanol for 15 minutes at −20°C, washed once in PBS, incubated 5 minutes in PBS + 0.1% triton-x-100, blocked for 1 hour at room temperature with PBS/2% FCS, then incubated overnight at 4°C with primary antibodies diluted in PBS/2% FCS. The next day, coverslips or Ibidi wells were washed two times with PBS and incubated for 1 hour at room temperature with Alexa-Fluor secondary antibodies from Invitrogen (Donkey anti-Rabbit IgG cat# A32790, Donkey anti-mouse IgG cat#A32773) diluted 1:1000 in PBS/2% FCS. DAPI (Sigma cat#D9542) was used to stain DNA. To objectively measure intensity of nuclear NRF2 immunofluorescence, ROIs of the nuclei were produced from images of DAPI fluorescence using ImageJ. These outlines were then overlayed onto the corresponding images of NRF2 immunofluorescence and mean intensity within each cardiomyocyte nuclei was recorded. After background correction, measurements from treated cells were normalized to untreated control cells within the same experiment.

### Cell Viability Testing with Trypan Blue Exclusion Method [44]

MCF7 cells were plated in 1.5 mls media in 6-well plates. After incubation with or without 500µM AICAr for 24 hours, media was replaced with fresh media containing 2µM doxorubicin or vehicle. After 6 hours, doxorubicin was removed, and replaced with fresh media overnight. Next, cells were trypsinized and resuspended in fresh media. Equal volumes of cell suspension and 0.4% trypan blue were mixed and analyzed for cell viability using CellDrop (DeNovix).

### Statistical Analysis

Statistical analysis was performed using GraphPad Prism version 10 (GraphPad Software, Boston, MA, USA). All values were expressed as mean ± standard error of the mean (SEM) from a minimum of three independent experiments. One-way analysis of variance (ANOVA) with Tukey’s post hoc comparison was used for testing multiple groups with normally distributed data. Dunnett’s was used for normally distributed data following ANOVA when comparing multiple treatment groups to a single control group. Data from multiple groups not normally distributed were analyzed using Kruskal-Wallis One Way Analysis of Variance on Ranks with Dunn’s Method for multiple pairwise comparisons. Student’s two-tailed T-test was used to determine significance between two groups, with Mann–Whitney rank sum test for data not normally distributed. Statistical significance was defined as P < 0.05. In Box and Whisker plots, whiskers denote the 5^th^-95^th^ percentile of the data.

## Results

### AICAr activates AMPK signaling but inhibits autophagy

To examine AICAr effects on AMPK signaling and autophagic flux, NRVMs were treated with increasing amounts of AICAr for 3 hours in the presence or absence of bafilomycin. As expected, AICAr dose-dependently increased AMPK^Thr172^ phosphorylation.

Consistent with increased AMPK activity, phosphorylation of AMPK substrate and autophagy regulator ULK^Thr555^ was also elevated by AICAr treatment. Surprisingly, however, the increase in these pro-autophagic signaling pathways was accompanied by a reduction in lipidated LC3 (LC3-II) and accumulation of autophagy substrate p62 SQST1. Co-treatment with bafilomycin, to inhibit lysosomal degradation, showed that AICAr reduced LC3-II levels relative to bafilomycin treatment alone, indicating that AICAr treatment suppresses LC3 lipidation and decreases LC3-II flux. Additionally, the AICAr-induced increase in p62 was not further elevated by bafilomycin, indicating that AICAr treatment strongly inhibits autophagic flux in cardiomyocytes by diminishing autophagosome formation (**Figure 1**).

**Fig. 1.**
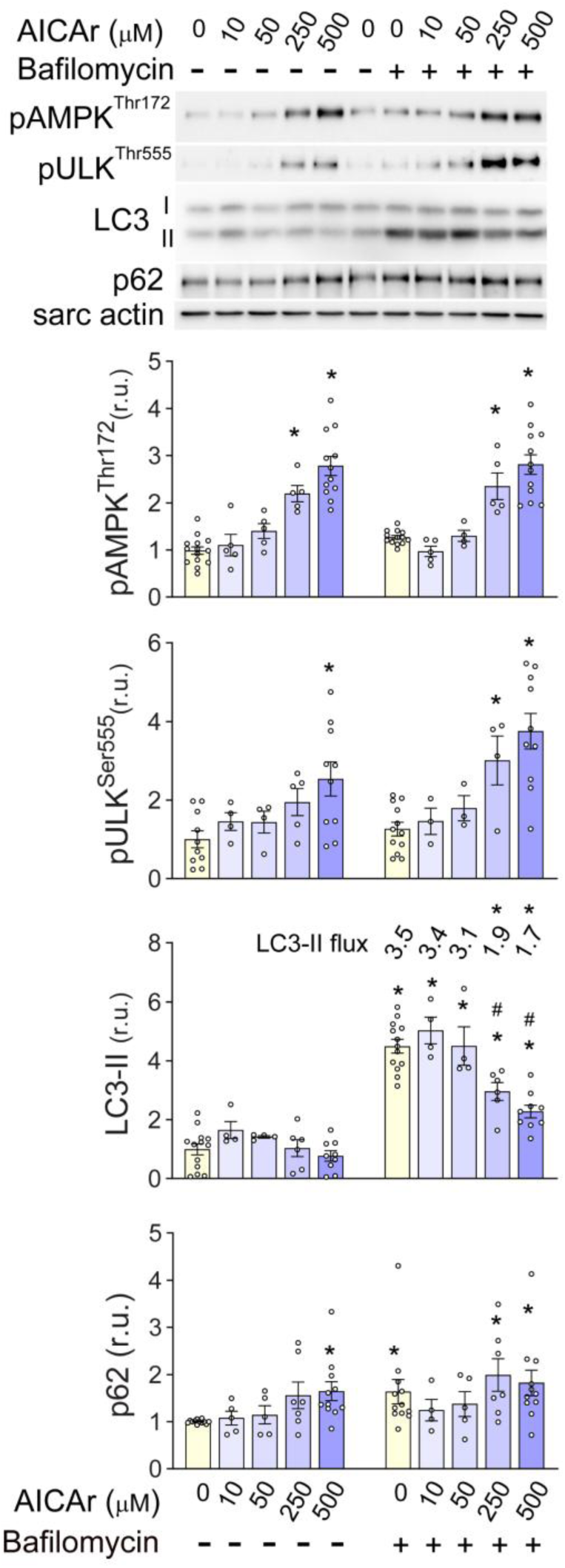
AICAr dose response effects on phosphorylation of AMPK, ULK1 and autophagic flux NRVMs were treated with 0, 10, 50, 250, or 500μM AICAr for 3 hours with or without 50nM bafilomycin for the last two hours. AMPKThr172 and UlkSer555 phosphorylation, LC3-II, autophagic flux, and p62 levels were analyzed by western blot and normalized to sarcomeric actin. * indicates p<0.05 vs control. # indicates p<0.05 comparing LC3-II control values and treatment groups under bafilomycin treatment

### AICAr-induced cardiomyocyte AMPK activation and autophagy inhibition is dependent upon ADK activity

To determine if autophagy inhibition by AICAr requires it’s phosphorylation by ADK, cells were treated with AICAr in the presence of ABT-702, an adenosine kinase inhibitor. HPLC analysis of ZMP levels in cardiomyocytes confirmed that ABT-702 completely prevented conversion of AICAr into ZMP (Supplemental figure 1).

Autophagic flux was examined by treating with bafilomycin for the last 2 hours of the experiment. ABT-702 blocked AICAr-induced phosphorylation of AMPK^Thr172^ and ULK^Ser555^, as well as the reduction in LC3-II and accumulation of p62 (**Figure 2a**), indicating that ZMP formation is required for both activation of AMPK and inhibition of autophagy.

**Fig. 2.**
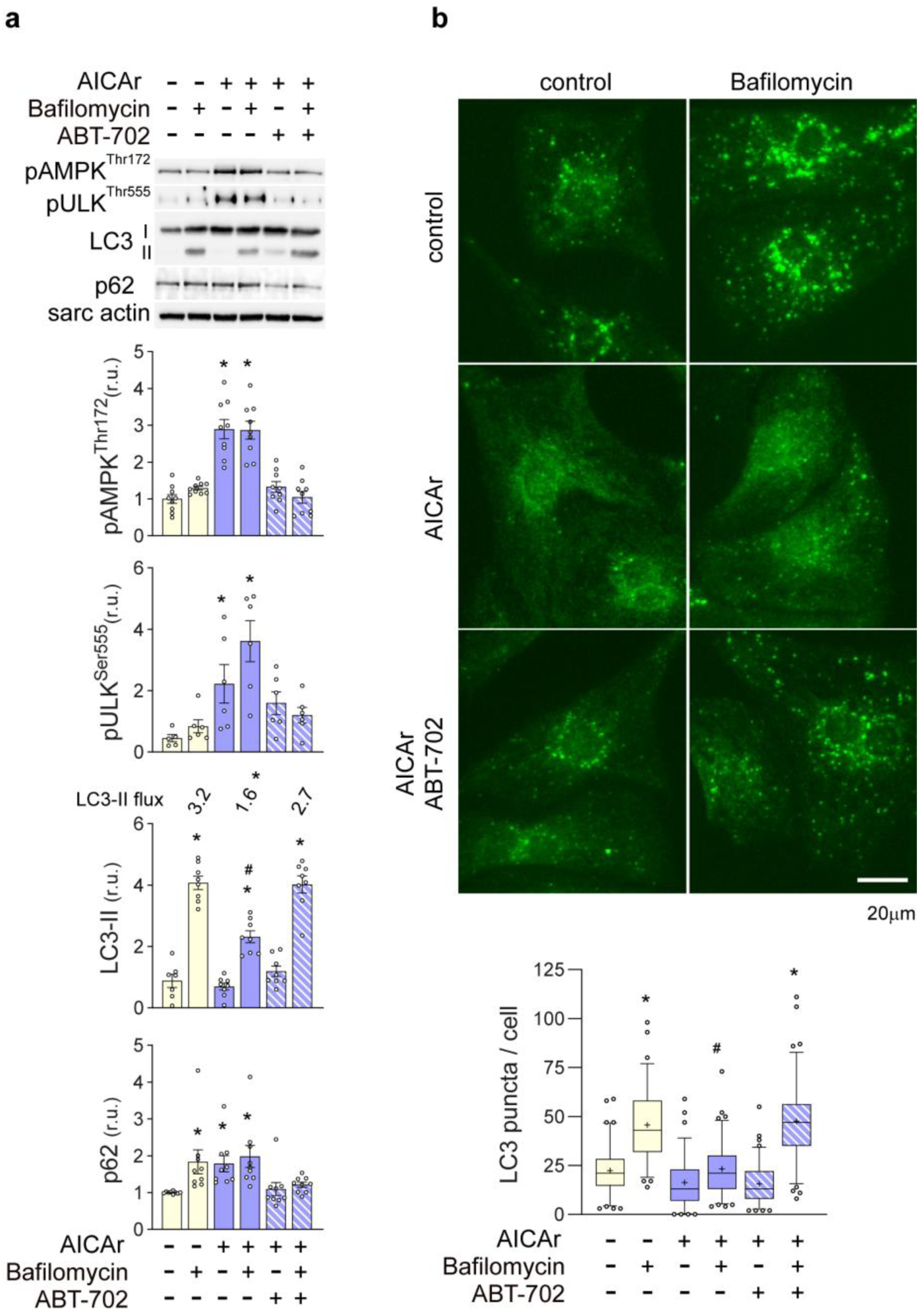
ADK inhibition with ABT-702 blocks AICAr suppression of autophagy **a** NRVMs were treated with 500μM AICAr with or without ABT-702 for 3 hours in the presence or absence of 50nM bafilomycin for the last 2 hours. AMPK^Thr172^, Ulk^Ser555^ phosphorylation, LC3-II, autophagic flux, and p62 levels were analyzed by western blot and normalized to sarcomeric actin. **b** In parallel experiments, LC3-positive puncta were examined by immunofluorescence. Box and Whisker plots represent the number of autophagosomes/cell from 3 independent experiments, counting 20-30 cells per condition in each experiment. The + sign indicates the mean. * indicates p<0.05 as compared to controls. # indicates p<0.05 comparing LC3-II control values and treatment groups under bafilomycin treatment

Immunofluorescence analysis also demonstrated that AICAr treatment diminishes formation of LC3-containing autophagosomes in the presence of bafilomycin, and this effect is blocked by ABT-702 (**Figure 2b**). Thus, the conversion of AICAr to ZMP by adenosine kinase mediates both activation of AMPK and inhibition of autophagy in NRVMs.

### AICAr inhibition of autophagy increases P62 and NRF2 signaling

To examine the longer-term effects of AICAr on cardiomyocyte autophagy inhibition and AMPK activation, a time-course study was performed. Autophagic flux was examined by adding 50nM bafilomycin for the last 2 hours of each time point (3, 6, and 24 hours) of AICAr treatment. As shown in (**Figure 3a**), phosphorylation

**Fig. 3.**
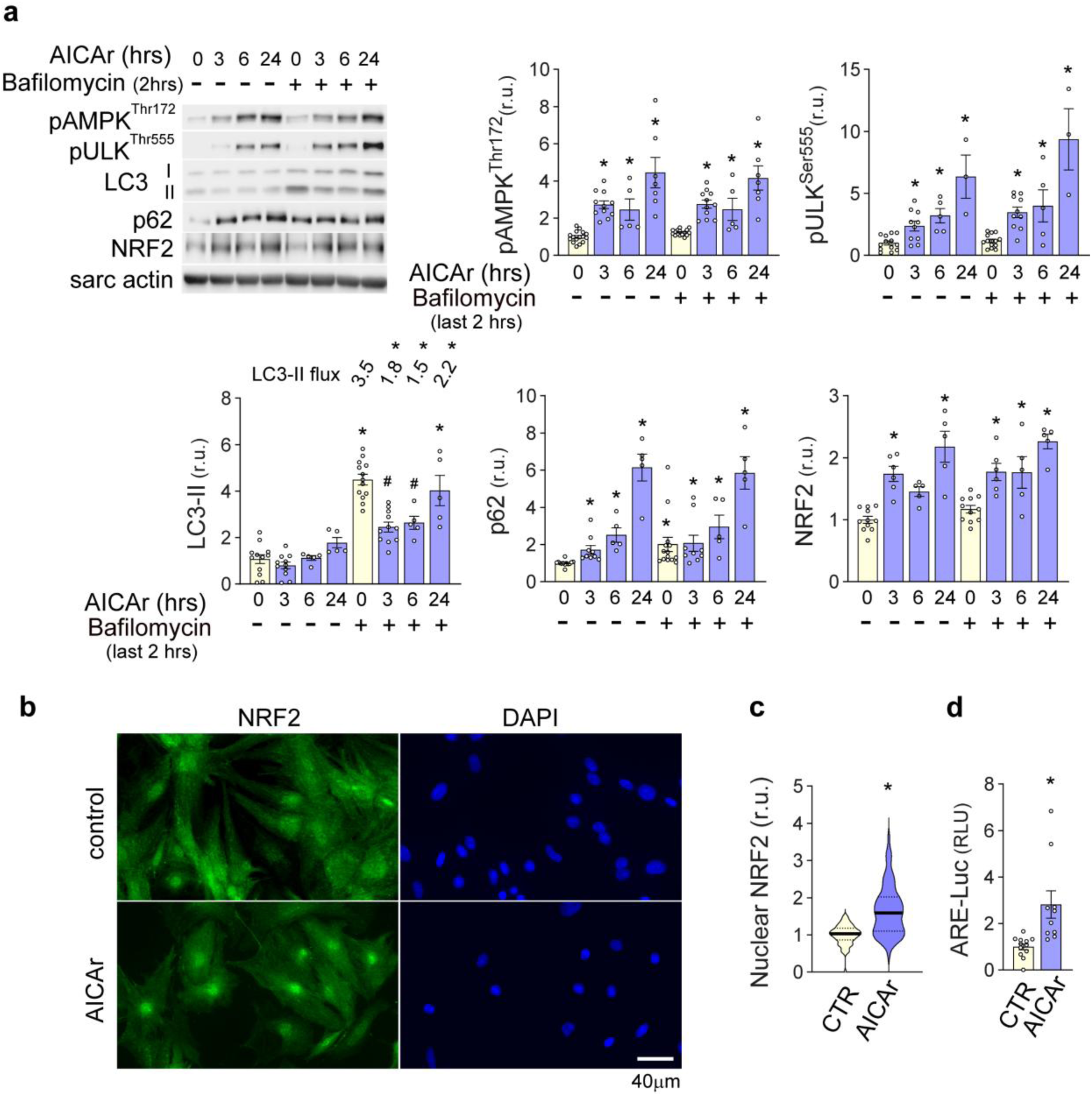
Time course of *AICAr effects on AMPK activation, autophagy, NRF2 expression and NRF2 transcriptional activity* **a** NRVMs were treated with 500µM AICAr for 3, 6, and 24 hours. To examine autophagic flux, 50nM bafilomycin was added for the last 2 hours. pAMPK^Thr172^, pUlk^Ser555^, p62, LC3-II, autophagic flux, and NRF2 levels were analyzed by western blot and normalized to sarcomeric actin. **b** immunofluorescent staining of NRF2 after 24 hours of AICAr treatment and **c** quantification of nuclear NRF2 from two separate experiments, each measuring 30-40 cells per condition. **d** ARE/NRF2 reporter luciferase activity was measured after 24 hours of AICAr treatment and normalized to co-transfected constitutive renilla luciferase activity. * indicates p<0.05 as compared to control. # indicates p<0.05 comparing LC3-II control values and treatment groups under bafilomycin treatment of AMPK^Thr172^ and ULK1^Ser555^ continued to increase through 24 hours. At the same time, LC3-II formation was diminished at 3 and 6 hours relative to bafilomycin treatment alone, but recovered nearly to control values by 24 hours. Autophagic flux remained significantly lower, however, suggesting autophagy remained impaired by AICAr. p62 also continued to accumulate in the presence of AICAr and did not increase further in the presence of bafilomycin. Because p62 is known to block KEAP1-dependent degradation of NRF2 [17], we examined whether AICAr-induced p62 accumulation could increase cardiomyocyte NRF2 protein levels. AICAr treatment increased NRF2 expression mostly in parallel with p62 accumulation, with peak expression occurring after 24 hours. At the same time, an increase in *p62* and *NRF2* mRNA levels was not observed until 24 hours (**Supplemental figure 2**), suggesting the increase in p62 and NRF2 protein at 3 and 6 hours occurs post-transcriptionally, while the increase in p62 and NRF2 protein at 24 hours also involves transcriptional upregulation. Nuclear translocation of NRF2 is important for inducing anti-oxidant gene expression. AICAr treatment increased both NRF2 nuclear localization (**Figure 3b** and **3c**), and activity of an anti-oxidant response element (ARE)-driven luciferase reporter (ARE-luciferase) (**Figure 3d**), indicating that AICAr induces NRF2 expression, nuclear localization and transcriptional activity.

### AICAr treatment protects against doxorubicin-induced ROS and cell death

Activation of NRF2 signaling has been shown to attenuate doxorubicin cardiotoxicity [28, 30, 52]. Therefore, we examined whether AICAr-induction of NRF2 could exert similar protection in NRVMs. Because oxidative stress is believed to contribute to doxorubicin cardiotoxicity [2], and NRF2 induces anti-oxidant gene expression [25], we first examined the effect of 24-hour AICAr pretreatment on doxorubicin-induced ROS formation in NRVMs.

As shown by DCFDA assay (**Figure 4a**), doxorubicin treatment significantly increased ROS, but this effect was diminished by AICAr pretreatment. We next examined whether AICAr pretreatment could preserve viability of NRVMs exposed to doxorubicin. An initial dose response study found that treatment of NRVMs with 2μM doxorubicin treatment for 6 hours reduced cell viability by approximately 50% when assayed 24 hours later by XTT assay (**Supplemental figure 3a**). Pretreatment of NRVMs with AICAr for 24 hours, however, significantly attenuated doxorubicin-induced loss of cell viability (**Figure 4b**). Preliminary studies also suggested 6hr pretreatment may also offer some protection against doxorubicin toxicity, but not to the same extent as 24 hours (**Supplemental figure 3a)**. These data indicate that AICAr pretreatment protects against doxorubicin-induced oxidative stress and injury in cardiomyocytes.

**Fig. 4.**
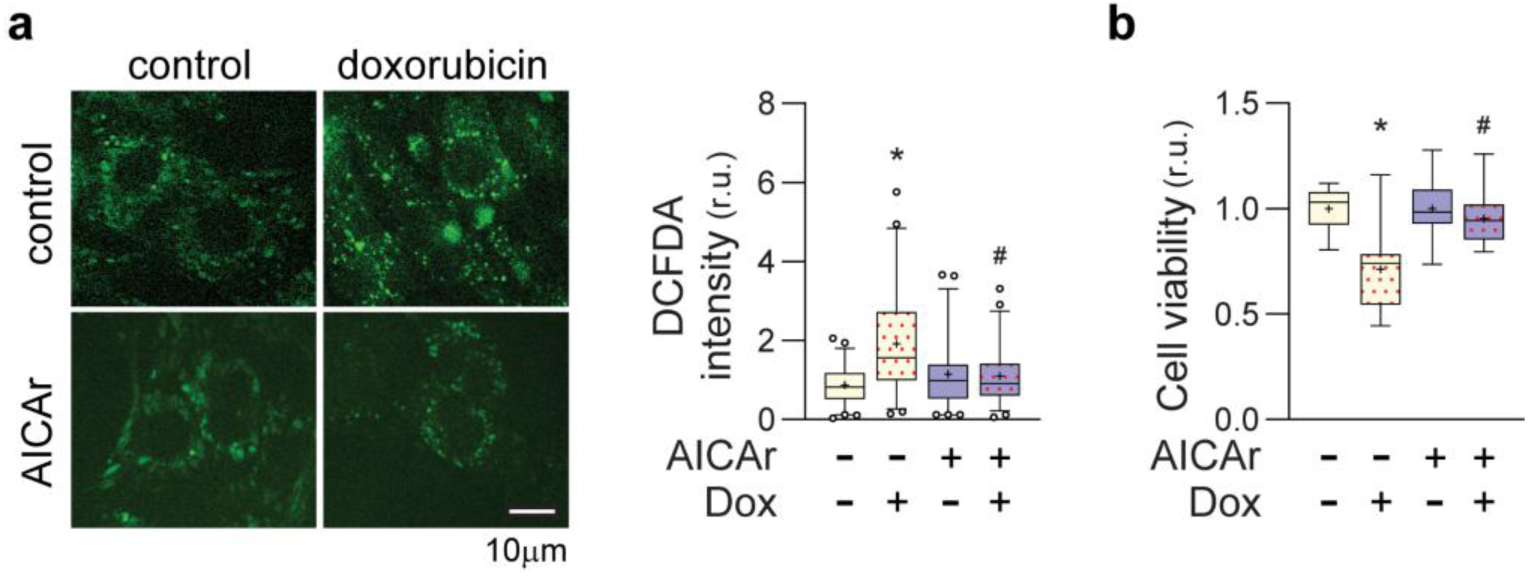
Effects of AICAr pretreatment on ROS and cell viability in NRVMs treated with doxorubicin **a** NRVM’s were pretreated with 500μM AICAr for 24 hours followed by 6 hours exposure to 2μM doxorubicin. ROS were measured via DCFDA assay and fluorescence microscopy. Box and Whisker plots represent data from 3 independent experiments. 20 cells from each condition were measured per experiment and normalized to intra-experiment controls. **b** NRVM’s were pretreated with 500μM AICAr for 24 hours prior to 6hrs exposure to 2μM doxorubicin. Cell viability was measured 24 hours later by XTT assay. Box and Whisker plots represent data from 3 independent experiments, each including 3-6 wells per condition. * indicates p<0.05 relative to control, # indicates p<0.05 comparing doxorubicin treatment to doxorubicin + AICAr treatment

### AMPK activation in absence of autophagy inhibition does not increase cardiomyocyte p62 and NRF2 expression

While many reports indicate that AMPK promotes autophagy, a recent study suggested that AMPK can inhibit autophagy [38]. Because phosphorylation of AMPKThr172, as well as the autophagy substrate p62, both increase through 24 hours of AICAr treatment, we examined the role of AMPK in cardiomyocyte p62 and NRF2 expression via adenoviral over-expression of AMPKα2. As shown in (**Figure 5**), AMPK^Thr172^ phosphorylation was significantly elevated by AMPKα2 over-expression, but this was associated with reduced p62 expression. NRF2 expression also trended lower as AMPK levels increased. These results suggest that AICAr-induced accumulation of p62 and NRF2 occurs despite AMPK activation, rather than because of it.

**Fig. 5.**
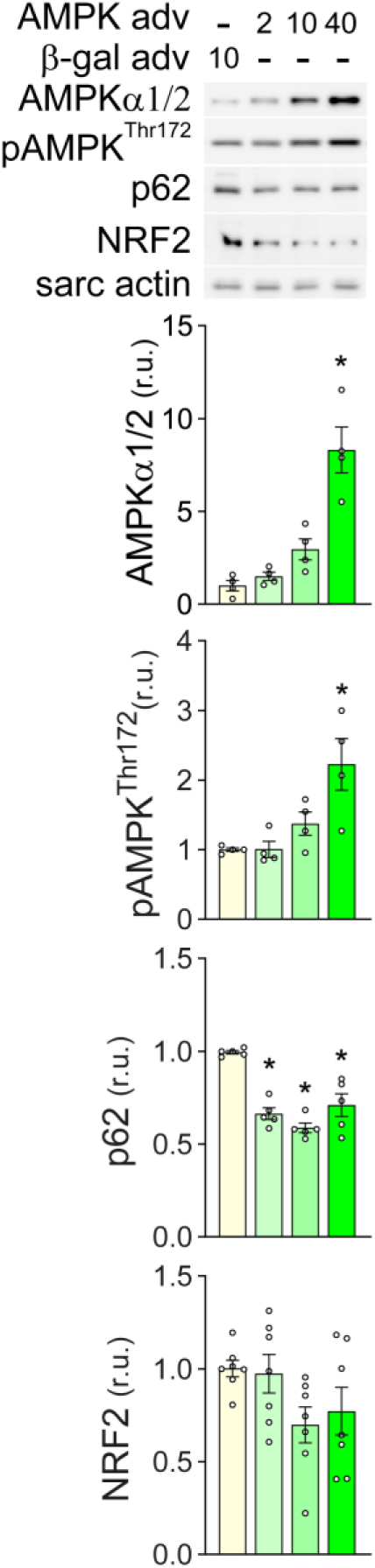
Effects of AMPKα2 over-expression on p62 and NRF2 in NRVMs Total AMPKα, pAMPK^Thr172^, p62, and NRF2 levels were examined by western blot 48 hours after infection with the indicated MOI of β-gal or wild type AMPKα2 adenovirus. * indicates p<0.05 relative to control

### AICAr-induced increases of NRF2, HO-1, and NQO1 gene expression and protection against doxorubicin-induced cardiotoxicity are dependent upon p62

To examine the role of p62 in AICAr-induced NRF2 expression, we depleted p62 using RNAi. As shown in (**Figure 6a**), p62 RNAi treatment dramatically lowered p62 as compared to cells treated with non-targeting control RNAi (CTR RNAi), but did not significantly reduce AMPK^Thr172^ phosphorylation. While AICAr significantly increased p62 and NRF2 protein content in CTR RNAi-treated cells, these increases were absent in p62 deficient cells, indicating p62 is necessary for AICAr-induced increase in NRF2 expression. To further examine the impact of p62 on AICAr-induced NRF2 and antioxidant gene expression, we examined the effects of p62 knockdown on mRNA levels of *p62, NRF2*, and NRF2 target genes *HO-1, NQO1, ATG7, and SOD1*. In control RNAi-treated cells, AICAr treatment significantly increased *p62, NRF2, HO-1, NQO1*, *SOD1* and *ATG7* mRNAs. p62 RNAi lowered *p62* mRNA by 90%. While p62 depletion did not affect basal mRNA levels of *NRF2, HO-1, NQO1*, *SOD1*, or *ATG7*, the AICAr-induced increases in *NRF2, HO-1, NQO1* and *ATG7* were eliminated (**Figure 6b**). Interestingly, the AICAr-induced increase in *SOD1* mRNA was not affected by p62 depletion, indicating AICAr can induce *SOD1* transcription through a p62/NRF2-independent mechanism. Supporting an important role for p62 in AICAr-induced NRF2 activity, nuclear localization of NRF2 in response to AICAr appeared much diminished in comparison to AICAr-treated control RNAi cells (**Figure 6c**). Consistent with a role for p62 in AICAr protection against doxorubicin injury, AICAr-induced doxorubicin tolerance was diminished in p62 RNAi-treated cells as compared to ctr RNAi-treated cells (**Figure 6d**). These data indicate that AICAr treatment increases expression of NRF2 and antioxidant genes and protects against doxorubicin toxicity through a p62-dependent mechanism.

**Fig. 6.**
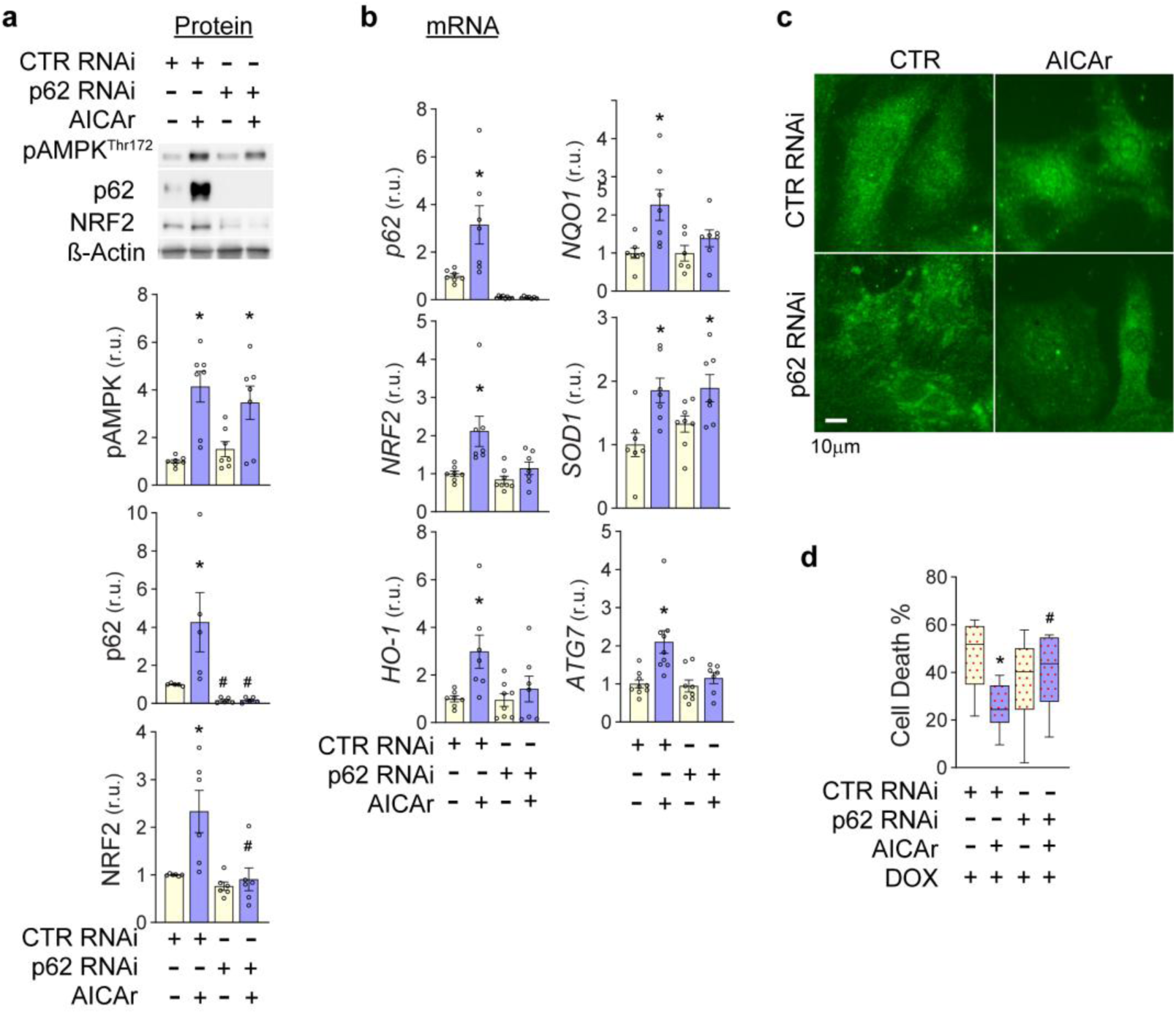
AICAr-induced increases of NRF2, NRF2 target genes, and protection against doxorubicin-induced cell death require p62 NRVMs were treated with control RNAi or p62 RNAi and exposed to 24 hours AICAr or control conditions. **a** pAMPK^Thr172^, p62 and NRF2 protein levels were measured by western blot and normalized to β-actin. **b** mRNA levels of *p62*, *NRF2*, *HO-1*, *NQO1*, *ATG7* and *SOD1* were measured by RT-QPCR and normalized to 18s RNA. **c** NRF2 localization was examined by immunofluorescence **d** Following 24 hours of AICAr treatment, control RNAi or p62 RNAi-treated NRVMs were exposed to 2µM doxorubicin for 6 hours. 24 hours later, % cell death was determined by XTT assay. Box and Whisker plots represent data from 3 independent experiments with 3-6 wells measured per condition and normalized to intra-experiment controls. * Indicates p<0.05 relative to ctr RNAi. # indicates p<0.05 comparing p62 RNAi to ctr RNAi exposed to the same treatments

### AICAr pretreatment preserves p62 and nuclear NRF2 during doxorubicin treatment

Oxidative stress is known to diminish KEAP-mediated degradation of NRF2 [51]. Doxorubicin induces oxidative stress [2] and has also been shown to either enhance [22] or diminish [43] cardiomyocyte autophagy. However, we did not observe a significant change in LC3-II flux of NRVM’s treated with doxorubicin for 3 or 6 hours (**Supplemental figure 4**). To further define how AICAr-induced p62/NRF2 expression countered doxorubicin effects on cardiomyocytes, p62 and NRF2 expression were examined after 6 hours of doxorubicin exposure with or without AICAr pretreatment. Doxorubicin treatment significantly decreased p62 protein levels but did not lower NRF2 expression (**Figure 7a**). In AICAr-pretreated cells, p62 and NRF2 remained elevated 6 hours after removing AICAr. In response to doxorubicin treatment, p62 protein levels decreased relative to AICAr-treated cells, but did not drop below untreated control values. Likewise, the elevated NRF2 expression observed in AICAr-treated cells appeared to decrease slightly after doxorubicin treatment, but still trended higher on average than control values. Because nuclear localization of NRF2 is necessary for ARE-dependent gene transcription [3], we examined the effect of doxorubicin and AICAr pretreatment on NRF2 nuclear localization. Even though treatment of NRVMs with 2μM Dox did not decrease overall NRF2 protein levels (**Figure 7a**), it significantly decreased NRF2 nuclear localization (**Figure 7b**) and (larger field; **Supplemental figure 5**). Importantly, pretreatment with AICAr increased nuclear localization of NRF2 and partially preserved nuclear NRF2 levels after exposure to doxorubicin. These findings suggest that doxorubicin treatment, in addition to inducing oxidative stress, may also lower the ability of cardiomyocytes to defend against oxidative stress by diminishing p62 and nuclear NRF2. Thus, a surplus of p62 and nuclear NRF2 induced by AICAr pretreatment may help preserve p62/NRF2 signaling in the presence of doxorubicin.

**Fig. 7.**
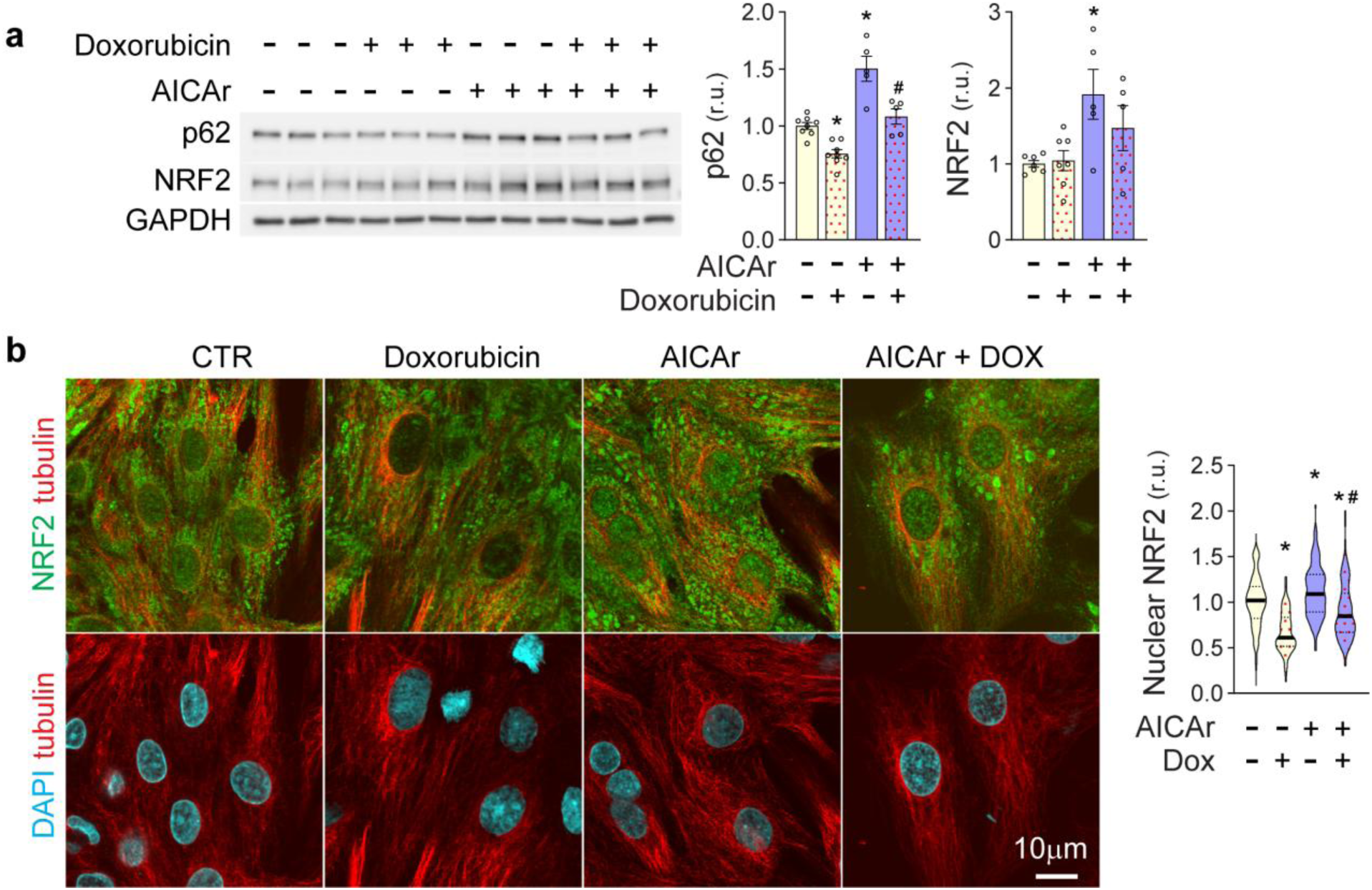
Doxorubicin effects on cardiomyocyte p62, NRF2, and Nuclear NRF2 After 24-hours pre-treatment with AICAr or control conditions, NRVMs were treated with 2µM doxorubicin for 6 hours. **a** P62 and NRF2 levels were analyzed by Western blot and normalized to GAPDH. **b** Nuclear NRF2 localization was measured by immunofluorescence analysis. Violin plots represent intensity of nuclear NRF2 stain from three independent experiments counting at least 40 nuclei per experiment, which were normalized to intra-experimental control values. * Indicates p<0.05 in comparison to control. # indicates p<0.05 comparing effects of doxorubicin to doxorubicin + AICAr.

### AICAr pretreatment does not induce p62/NRF2 and enhances doxorubicin chemotherapeutic efficacy in the MCF7 breast cancer cell line

An important property of any preventative against doxorubicin cardiotoxicity is that it does not diminish chemotherapeutic efficacy of doxorubicin in tumor cells. Since ADK appears necessary for increases of both AMPK activity and p62 by AICAr in NRVMs, we compared ADK expression in NRVMs, mouse hearts and the breast cancer cell line, MCF7. As shown in (**Figure 8A**), ADK levels are significantly lower in MCF7 cells than in the mouse heart or NRVMs. While NRVMs express predominantly the nuclear form of ADK (ADK-L; ∼40kD MW), and mouse hearts express both ADK-L and the cytosolic isoform (ADK-S; ∼38kD MW), MCF7 cells predominantly express ADK-S. Consistent with an important role for ADK in AICAr inhibition of autophagy, autophagic flux of MCF7 cells was only slightly diminished after 3 hours (**Figure 8B**). While there was a modest but significant decrease in autophagic flux after 24 hours of AICAr treatment, this appeared to result from impaired MCF7 cell degradation of LC3-II, rather than reduced LC3 lipidation, which we observed in NRVMs. (**Figure 8C**). While AMPK^Thr172^ phosphorylation levels trended higher in MCF7 cells in response to 24 hours of AICAr treatment, neither p62 or NRF2 were significantly increased after 3 or 24 hours of AICAr treatment, indicating that AICAr does not activate p62/NRF2 signaling in MCF7 cells.

**Fig. 8.**
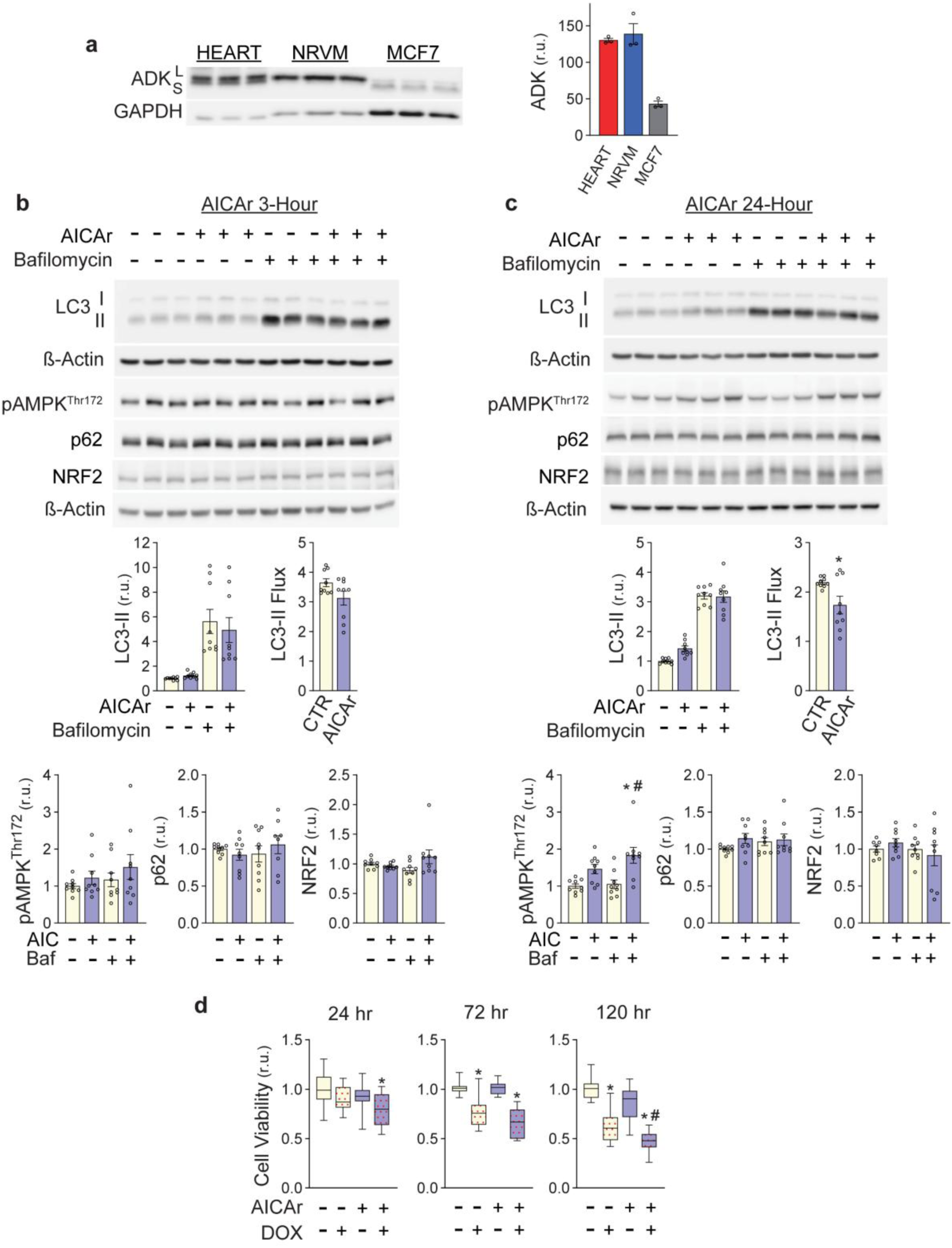
AICAr effects on autophagy, p62, NRF2 and doxorubicin toxicity in MCF7 cells **a** Western analysis of ADK in 20µg protein lysates from mouse hearts, NRVMs, and MCF7 cells. ADK-L and S are indicated. **b** MCF7 cells were treated with 500µM AICAr for 3 hours or **c** 24 hours. 50nM bafilomycin was added for the last 2 hours. LC3-II, LC3-II Flux, AMPK^Thr172^ phosphorylation, p62 and NRF2 levels were analyzed by Western blot and normalized to ß-Actin. **d** MCF7 cells were incubated with 500µM AICAr or control conditions for 24 hours, then exposed to 2µM doxorubicin for 6 hours. Cell viability was measured via XTT assay 24, 72, or 120 hours after doxorubicin treatment. Box and Whiskers plots represent data from 3 independent experiments. 4-5 wells per condition were examined in each experiment and normalized to intra-experiment controls. * Indicates p<0.05 relative to control, # indicates p<0.05 for doxorubicin treatment compared to doxorubicin + AICAr treatment

To find out if AICAr pretreatment interferes with chemotherapeutic effects of doxorubicin, MCF7 cell viability was measured by XTT assay. Doxorubicin decreased MCF7 cell viability more slowly than in NRVMs, as a significant reduction in cell viability was not observed until 5 days after doxorubicin removal (**Figure 8d**). Pretreatment with AICAr did not significantly affect cell viability on its own, but did accelerate and exacerbate doxorubicin-induced loss of MCF7 cell viability (**Figure 8d**). Analysis of cell viability using the Trypan Blue assay also indicated that pretreatment of MCF7 cells with AICAr sensitized MCF7 cells to doxorubicin **(Supplemental figure 6)**. Interestingly, unlike cardiomyocytes, many MCF7 cells became unattached 24 hours after treatment with doxorubicin or doxorubicin plus AICAr pretreatment (6.6 % of doxorubicin-treated and 9.4% of doxorubicin/AICAr-treated cells became detached, as compared to 0.73% of control or 1.4% of AICAr). To determine if these cells were still viable, media containing the detached cells was replated and maintained for 10 days, followed by Coomassie staining to assess new colony formation. Cells treated with doxorubicin or doxorubicin plus AICAr pre-treatment formed no new colonies, while control or AICAr-treated cells formed 70-80 new colonies. These data indicate that cells which detached after doxorubicin exposure are not viable. Thus, AICAr pretreatment does not interfere with and actually appears to enhance chemotherapeutic effects of doxorubicin on MCF7 cells (**Supplemental figure 6**).

## Discussion

This study identified a novel mechanism by which the widely used AMPK activating compound, AICAr, exerts a cytoprotective response in cardiomyocytes. Despite strongly activating AMPK, AICAr inhibits cardiomyocyte autophagy, leading to p62 accumulation, NRF2 upregulation, anti-oxidant gene expression and resistance to doxorubicin toxicity. Transient autophagy inhibition by AICAr and accumulation of p62 depends upon ADK activity, so that MCF7 breast cancer cells, which poorly express ADK, are not protected by AICAr. Instead, AICAr was found to sensitize MCF7 cells to doxorubicin-induced cell death. This raises the possibility that differences in ADK expression between the heart and breast cancer cells might be exploited by pretreating with AICAr or similar drugs to selectively protect cardiomyocytes against doxorubicin toxicity while enhancing its chemotherapeutic efficacy in tumors.

### Inhibition of autophagy by AICAr

While AICAr is a potent AMPK activator, it can also exert several AMPK-independent actions [50], including inhibition of autophagy. AICAr inhibition of autophagy was initially observed in hepatocytes [40] and more recently in skin fibroblasts [49] and pancreatic β-cells [15]. Here we show that AICAr, despite increasing phosphorylation of AMPK^Thr172^ and its downstream autophagy regulator, ULK1^Ser555^, is a potent inhibitor of autophagy in neonatal rat cardiomyocytes. The molecular mechanism(s) by which AICAr inhibits autophagy is unknown, but it is dependent upon adenosine kinase-dependent conversion of AICAr into ZMP (shown here and [40]) and appears to impede autophagy at the level of autophagosome initiation or development. Thus, while AICAr certainly elicits AMPK signaling in cultured NRVMs, our data suggest the AMPK-independent impact of autophagy inhibition should also be considered when assessing the physiological effects of this compound, particularly in cells or tissues that express high levels of ADK. Interestingly, a recent *in vivo* study in a rat model of doxorubicin cardiotoxicity also showed that AICAr pretreatment can attenuate doxorubicin cardiotoxicity even though an increase in cardiac AMPK phosphorylation was not detected. While the effects of AICAr on autophagy were not examined, this study supports the proposition that some cardioprotective actions of AICAr may be AMPK-independent [6].

Though prolonged impairment of autophagy is detrimental to cell fitness, our study demonstrates, for the first time, that transient autophagy inhibition by AICAr and subsequent p62-dependent induction of NRF2 signaling may be used as a preventative prior to doxorubicin treatment. The finding that autophagy inhibition can attenuate doxorubicin damage is partly supported by some previous studies [45, 46], where genetic or drug-induced autophagy inhibition during doxorubicin treatment was shown to be protective in cultured cells and *in vivo*. While protection against doxorubicin by autophagy inhibition could implicate excessive autophagy as a mechanism of doxorubicin cardiotoxicity, the role of autophagy in doxorubicin toxicity is complicated, as autophagy activation has also been shown to attenuate doxorubicin injury [43]. In our hands, doxorubicin had no effect on LC3-II turnover in NRVMs, but did lower p62 levels, which was associated with reduced nuclear NRF2. Because exogenous [26] or drug-induced [28, 52] NRF2 expression attenuates doxorubicin cardiotoxicity, the longer-lasting secondary effects of autophagy inhibition (*i.e.* activation of the p62/NRF2 positive feedback loop [16]), may provide an additional explanation for protection against cardiotoxicity by autophagy inhibitors.

### AICAr pretreatment induces at least two cardioprotective pathways

While our data indicate that autophagy inhibition and p62 accumulation play an important role in AICAr-induced protection, activation of AMPK by AICAr may also help shield against doxorubicin cardiotoxicity through previously identified mechanisms [47]. In addition, both AMPK and NRF2 have been implicated in reducing oxidative stress and ferroptosis and in preserving mitochondrial function, so that simultaneous increases of AMPK and NRF2 by AICAr may exert greater protective actions than activation of either pathway alone [39]. Direct interactions between components of the AMPK and the p62/NRF2 pathway have also been observed. For example, p62 can facilitate ULK1 and AMPK interaction, which promotes autophagic degradation of KEAP1 and subsequent NRF2 activity [24]. There is also evidence that AMPK phosphorylation of NRF2 can increase its nuclear localization [19]. AICAr-induction of p62/NRF2 via autophagy inhibition, along with simultaneous activation of AMPK, may thus synergistically protect against doxorubicin cardiotoxicity.

Interestingly, after 24 hours of AICAr treatment, LC3-II synthesis returned toward control values, suggesting a feedback mechanism may become engaged over time to overcome deficient LC3 lipidation. Besides increasing expression of anti-oxidant genes, NRF2 can directly promote transcription of several other autophagy genes besides p62, including ULK1, ATG5, ATG7, GABARAPL1 [36], and TFEB [34], a master transcriptional regulator of autophagy. Indeed, we observed an increase in ATG7 mRNA 24 hours after AICAr treatment that was dependent upon p62. ATG7 is the E1 enzyme that activates LC3 and LC3-related proteins that are critical for proper autophagosome development. The p62/NRF2-dependent increase in expression of ATG7 and other autophagy-related genes may thus help explain how prolonged treatment enhances autophagy in some models, even though it acutely inhibits autophagy. AMPK also promotes transcriptional regulation of autophagy by phosphorylating TFEB [37]. Thus, in addition to inducing NRF2-dependent anti-oxidant response, activation of AMPK and upregulation of NRF2 may help amass cellular autophagy and lysosomal machinery to facilitate disposal of damaged proteins or mitochondria associated with doxorubicin toxicity [4].

### AICAr sensitizes MCF7 cells to doxorubicin toxicity

Our data indicates that AICAr pretreatment not only protects cardiomyocytes but also enhances doxorubicin effects on MCF7 cells, suggesting potential dual benefit for doxorubicin chemotherapy. There is also evidence that AICAr can independently slow cell growth in some types of cancers [1, 5]. While we did not observe a significant effect of AICAr alone on MCF7 cell viability, our findings are consistent with other studies showing that AICAr can sensitize cancer cells to chemotherapeutics, including doxorubicin [11]. The mechanism(s) by which AICAr sensitizes MCF7 cells to doxorubicin remain unresolved, as both AMPK-dependent [11] and independent [1] mechanisms have been implicated. In our hands, AMPK^Thr172^ phosphorylation, an indicator of AMPK activation, did increase significantly after 24 hours of AICAr treatment but did not increase to the same extent as in NRVMs. At the same time, a slight reduction in autophagic flux was observed, but this appeared to occur at the level of LC3-II degradation, rather than lipidation, as found in NRVMs.

The poor activation of AMPK by AICAr and the minor effect of AICAr on autophagy, p62, and NRF2 in MCF7 cells could be attributed to lower levels of ADK expression in MCF7 cells as compared to NRVMs. In contrast to MCF7 cells, AICAr does induce NRF2 expression in hepatocellular carcinoma cells via an ADK-dependent, AMPK-independent mechanism [41]. Hepatocellular carcinoma cells are derived from hepatocytes, which express extremely high levels of ADK. Thus, while AICAr pretreatment shields cardiomyocytes and sensitizes MCF7 cells to doxorubicin, the impact of AICAr on doxorubicin chemotherapeutic efficacy in other cancers will likely vary, depending upon ADK expression level, metabolic routes of AICAr or ZMP, and pre-existing p62/NRF2 expression in the targeted cancer cell type.

### AICAr use in humans

AICAr has been investigated in a small phase I/II clinical trial for treatment of relapsed chronic lymphocytic leukemia (CLL) [48] and in larger clinical trials as a potentially cardioprotective agent during coronary artery bypass surgery [31, 33]. Interestingly, the maximum tolerated dose in the leukemia study (210mg/kg) resulted in ∼0.9mM concentration in whole blood after 4 hours of infusion, which is nearly twice the concentration of AICAr used in our study to inhibit autophagy and induce p62/NRF2 in cardiomyocytes. While the plasma half-life of AICAr in humans is only ∼1 hour [48], a significant portion of AICAr is taken up into cells and converted into ZMP. The extent to which ZMP accumulates and turns over in cardiomyocytes under a given dose will likely be the main determinant of its protective actions in the heart. It will thus be important to examine whether clinically tolerable doses of AICAr are sufficient to induce p62/NRF2 and cardioprotection in an *in vivo* model of doxorubicin toxicity.

## Summary

This study identified a novel protective mechanism of the widely used AMPK activator, AICAr. ADK-dependent phosphorylation of AICAr not only induces AMPK signaling but also inhibits cardiomyocyte autophagy, causing p62 accumulation, subsequent induction of NRF2 signaling and anti-oxidant gene expression. Transient autophagy inhibition by AICAr pretreatment thus helps shield cardiomyocytes against doxorubicin toxicity. Importantly, AICAr pretreatment does not protect breast cancer MCF7 cells and instead sensitizes them to doxorubicin. These opposite effects of AICAr on doxorubicin resistance in cardiomyocytes and breast cancer cells suggest that AICAr pretreatment has potential to provide dual benefits in doxorubicin chemotherapy.

## Statements and Declarations

The authors have no conflicts of interest to declare that are relevant to the content of this article. Data availability: Raw data will be made available upon request.

## Author contributions

JTF and EKF conceptualized the study. JTF and EKF wrote the manuscript. EKF and JTF designed experiments and collected data. JTF and BM reviewed and edited the manuscript. All authors have read and approved of the final manuscript. The authors would also like to thank Heike Stessel for excellent help with HPLC analysis.

## Supplemental data

**Supplemental Fig. 1.**
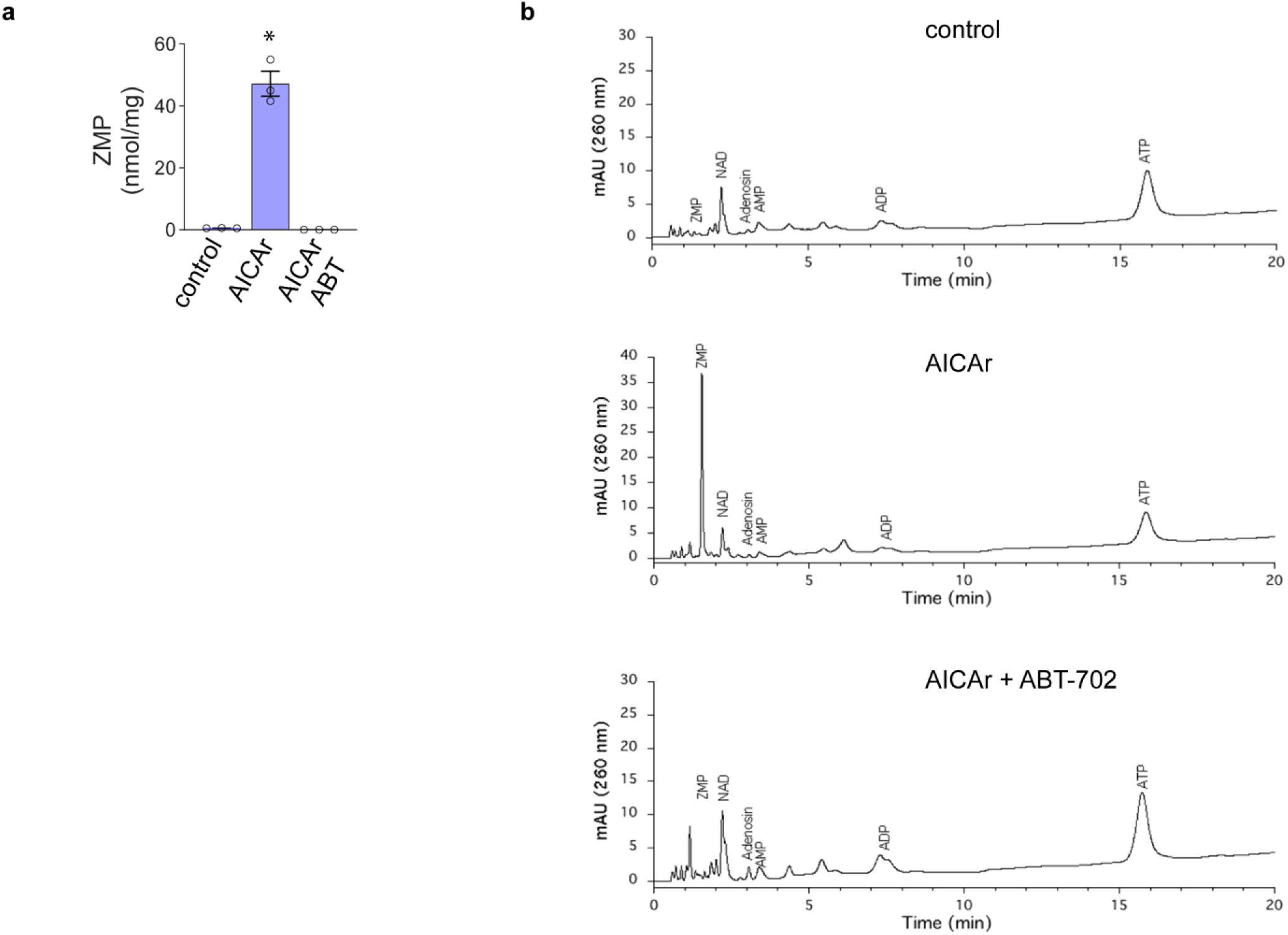
Effect of ADK inhibitor ABT-702 on ZMP formation in NRVMs NRVMs were treated for 3 hours with 500µM AICAr in presence or absence of 0.5 µM ADK inhibitor, ABT-702. ZMP levels were measured by HPLC and normalized to cellular protein content. Representative HPLC profiles are shown for the different treatment groups

**Supplemental Fig. 2.**
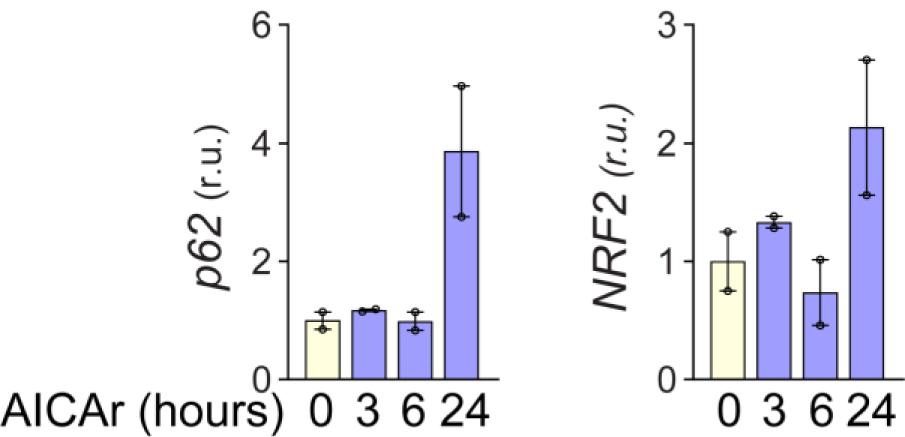
NRVM mRNA time course for p62 and NRF2 NRVMs were treated with 500μM AICAr for 3,6, and 24 hours prior to RNA collection. p62 and NRF2 mRNA was analyzed by RT-qPCR and normalized to 18S RNA. Results are expressed as mean +/- SEM. N=2 biologic replicates for each condition

**Supplemental Fig. 3.**
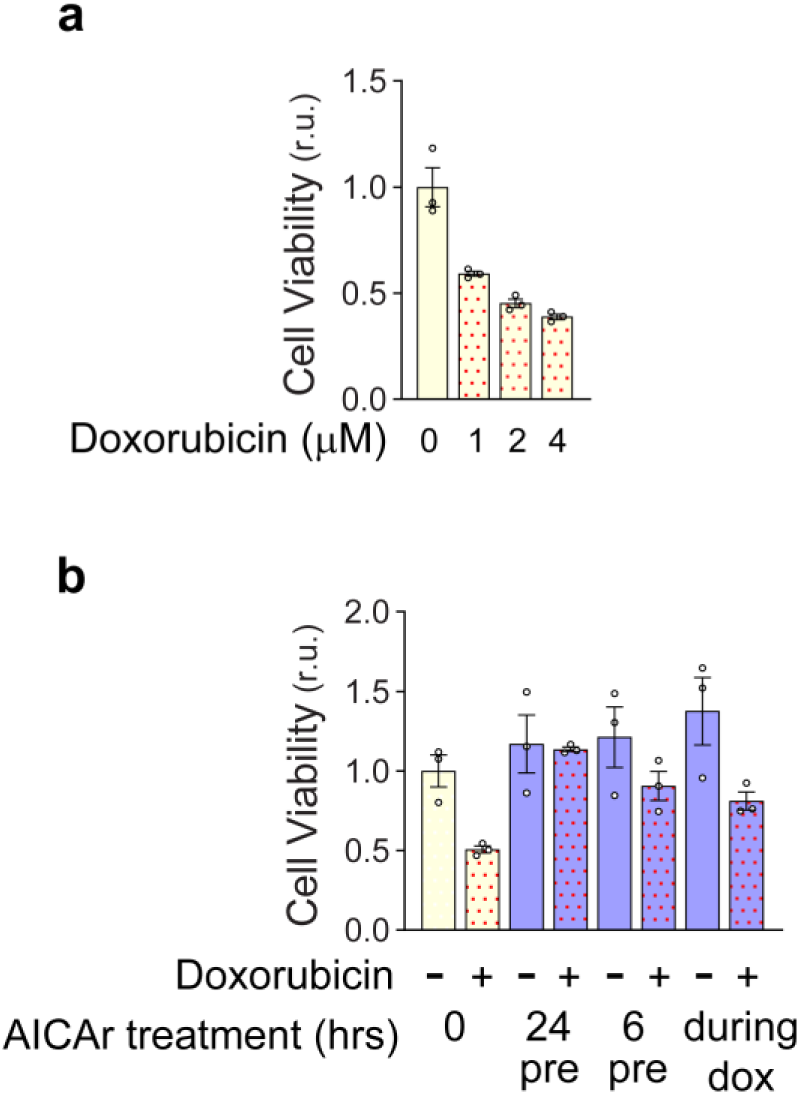
Doxorubicin dose response a NRVMs were treated with 0, 1, 2, or 4µM doxorubicin for 6 hours in triplicate. XTT assays were performed 24 hours after removal of doxorubicin. b NRVMs were pretreated with 500µM AICAr 24hrs prior, 6hrs prior, or during 6 hours exposure to 2µM doxorubicin. Results are expressed as mean +/- SEM

**Supplemental Fig. 4.**
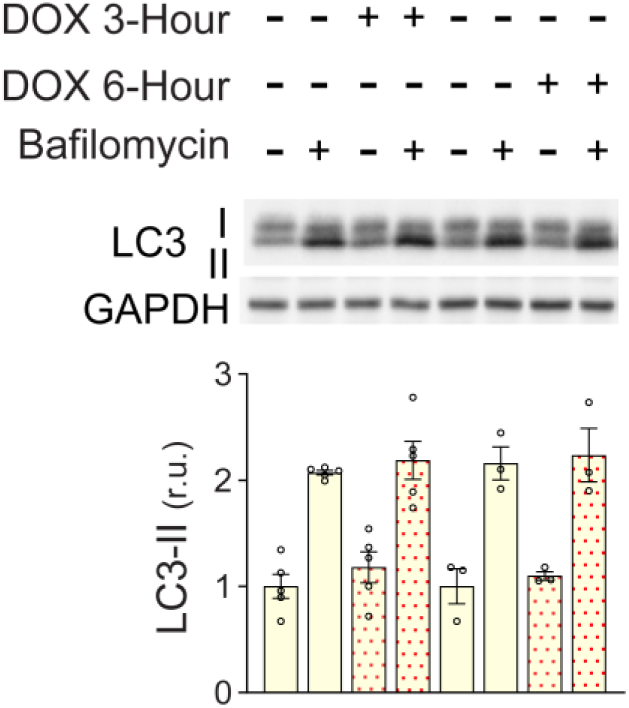
Doxorubicin effect on NRVM autophagy NRVMs were treated with 2μM doxorubicin for 3 or 6 hours in the presence or absence of 50nM bafilomycin for the last 2 hours. LC3 was analyzed via Western blot and normalized to GAPDH. Results are expressed as mean +/- SEM

**Supplemental Fig. 5.**
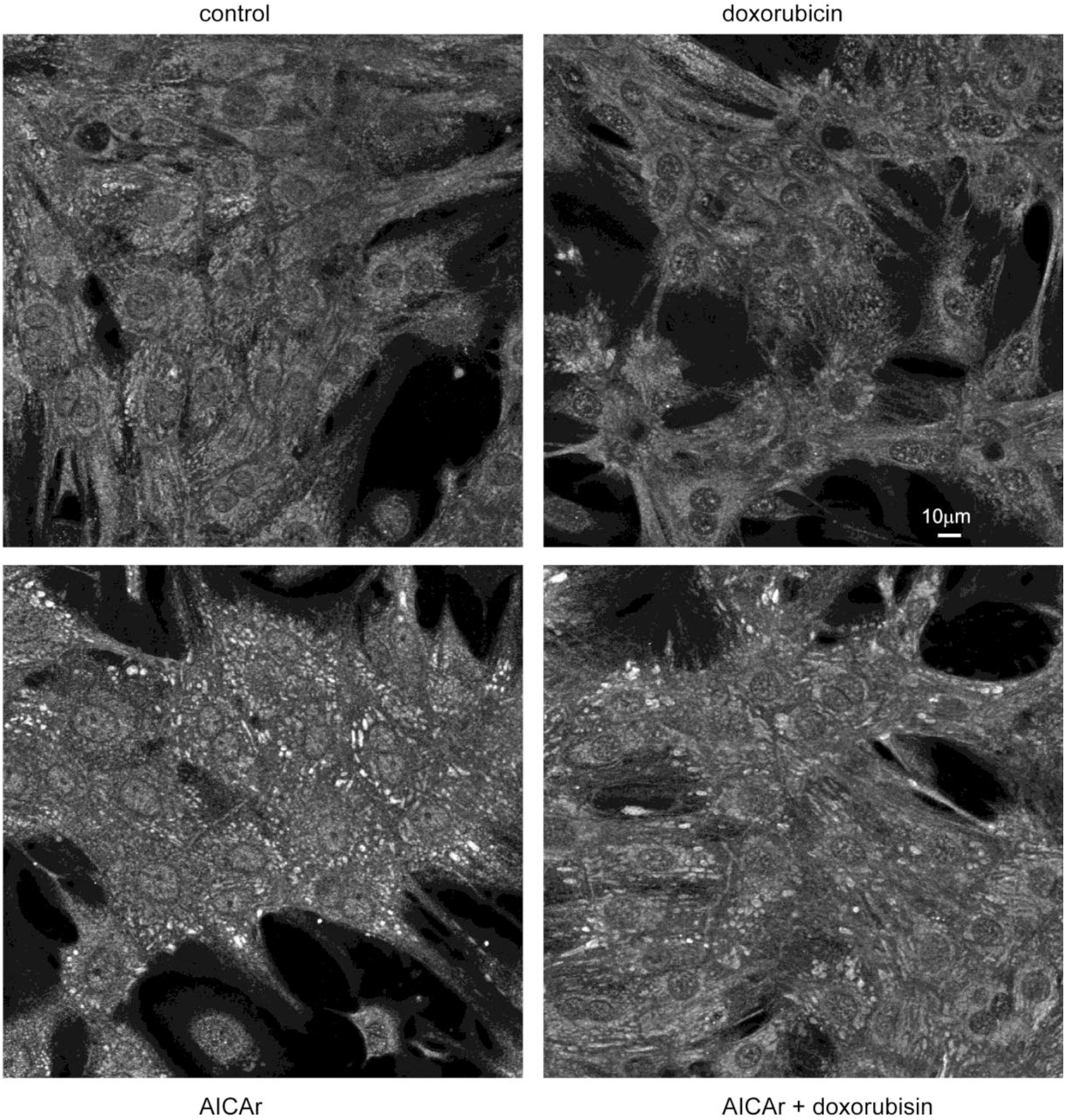
AICAr and doxorubicin effects on NRF2 nuclear localization NRF2 immunofluorescence of NRVMs pretreated for 24 hours with or without AICAr, followed by 6 hours of fresh media with or without doxorubicin.

**Supplemental Fig. 6.**
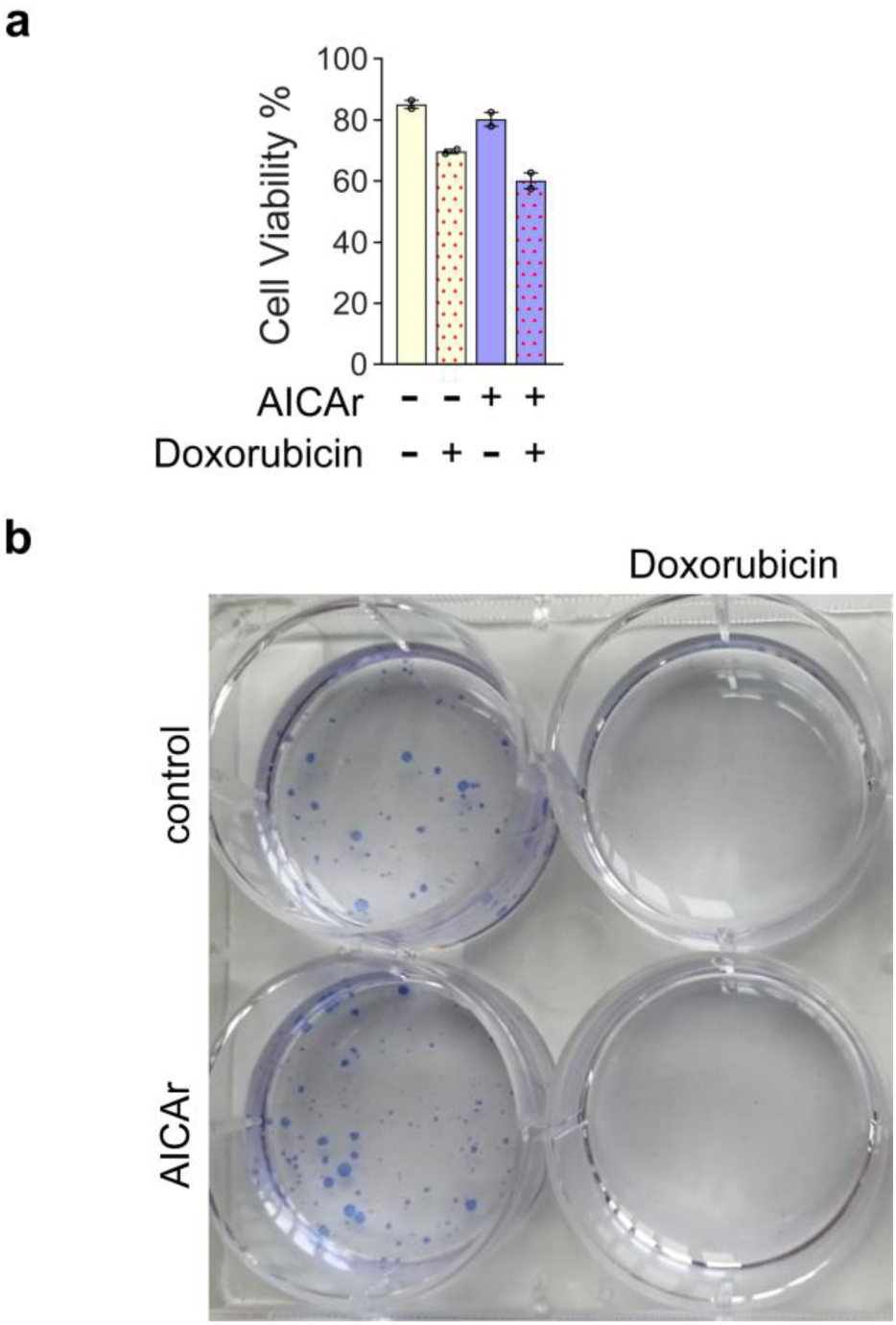
MCF7 Cells are not protected against Doxorubicin following pre-treatment with AICAr a MCF7 cells were cultured in the presence or absence of 500μM AICAr for 24 hours followed by 2μM doxorubicin for 6 hours. 24 hours later, Cell viability % (Live cells per ml/Total cells per ml) was analyzed via Trypan Blue Assay & CellDrop. Results are expressed as mean +/- SEM. * Indicates p<0.05 relative to control. N = 2 independent experiments. **b** The media containing floating cells was removed prior to the Trypan Blue Assay and replated onto a new 6-well plate. The plate was maintained for 10 additional days after which new cell colonies were identified by Coomassie staining

