## supplemental data for "AICAr inhibition of cardiomyocyte autophagy promotes p62-dependent NRF2 expression and protection against doxorubicin toxicity"

**Supplemental Figure 1**

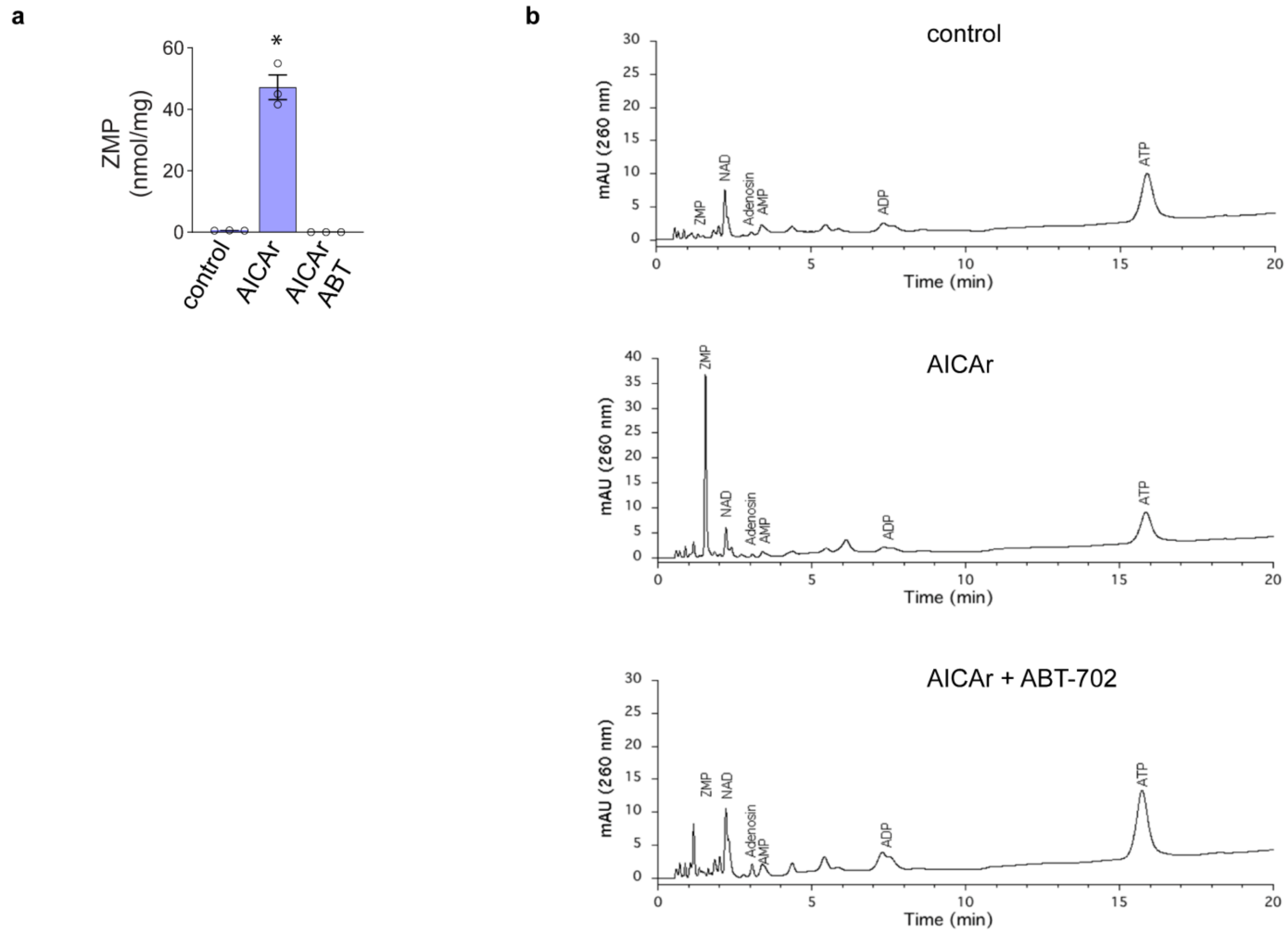


**Supplemental Fig. 1 Effect of ADK inhibitor ABT-702 on ZMP formation in NRVMs**

NRVMs were treated for 3 hours with 500μM AICAr in presence or absence of 0.5 μM ADK inhibitor, ABT-702. ZMP levels were measured by HPLC and normalized to cellular protein content. Representative HPLC profiles are shown for the different treatment groups

**Supplemental Fig. 2**

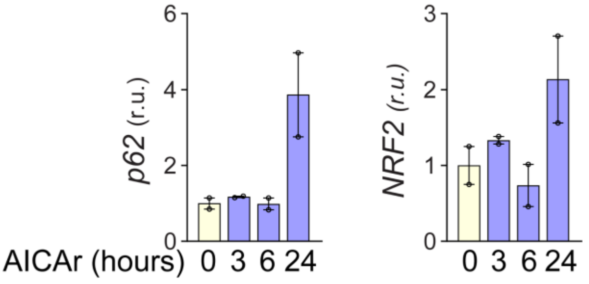


**Supplemental Fig. 2 NRVM mRNA time course for p62 and NRF2**
NRVMs were treated with 500µM AICAr for 3,6, and 24 hours prior to RNA collection. p62 and NRF2 mRNA was analyzed by RT-qPCR and normalized to 18S RNA. Results are expressed as mean +/- SEM. N=2 biologic replicates for each condition

**Supplemental Fig. 3**

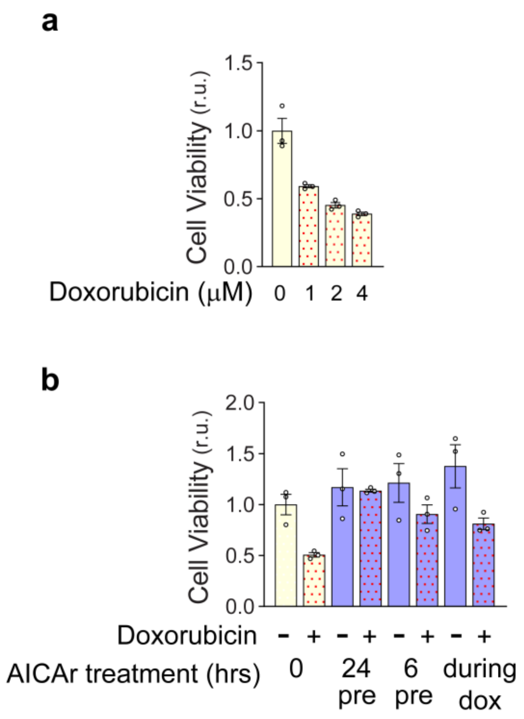


**Supplemental Fig. 3 Doxorubicin dose response**
**a** NRVMs were treated with 0, 1, 2, or 4µM doxorubicin for 6 hours in triplicate. XTT assays were performed 24 hours after removal of doxorubicin.
**b** NRVMs were pretreated with 500µM AICAr 24hrs prior, 6hrs prior, or during 6 hours exposure to 2µM doxorubicin. Results are expressed as mean +/- SEM

### Supplemental Fig. 4

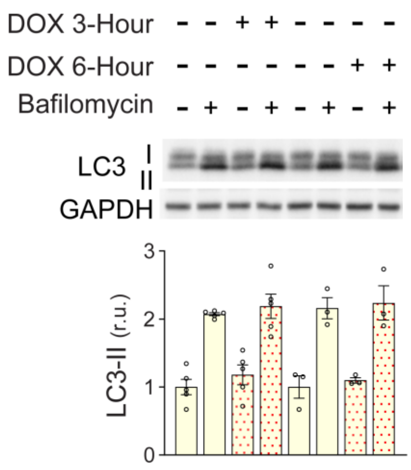


**Supplemental Fig. 4 Doxorubicin effect on NRVM autophagy**

NRVMs were treated with 2μM doxorubicin for 3 or 6 hours in the presence or absence of 50nM bafilomycin for the last 2 hours. LC3 was analyzed via Western blot and normalized to GAPDH. Results are expressed as mean +/- SEM

**Supplemental Fig. 5**

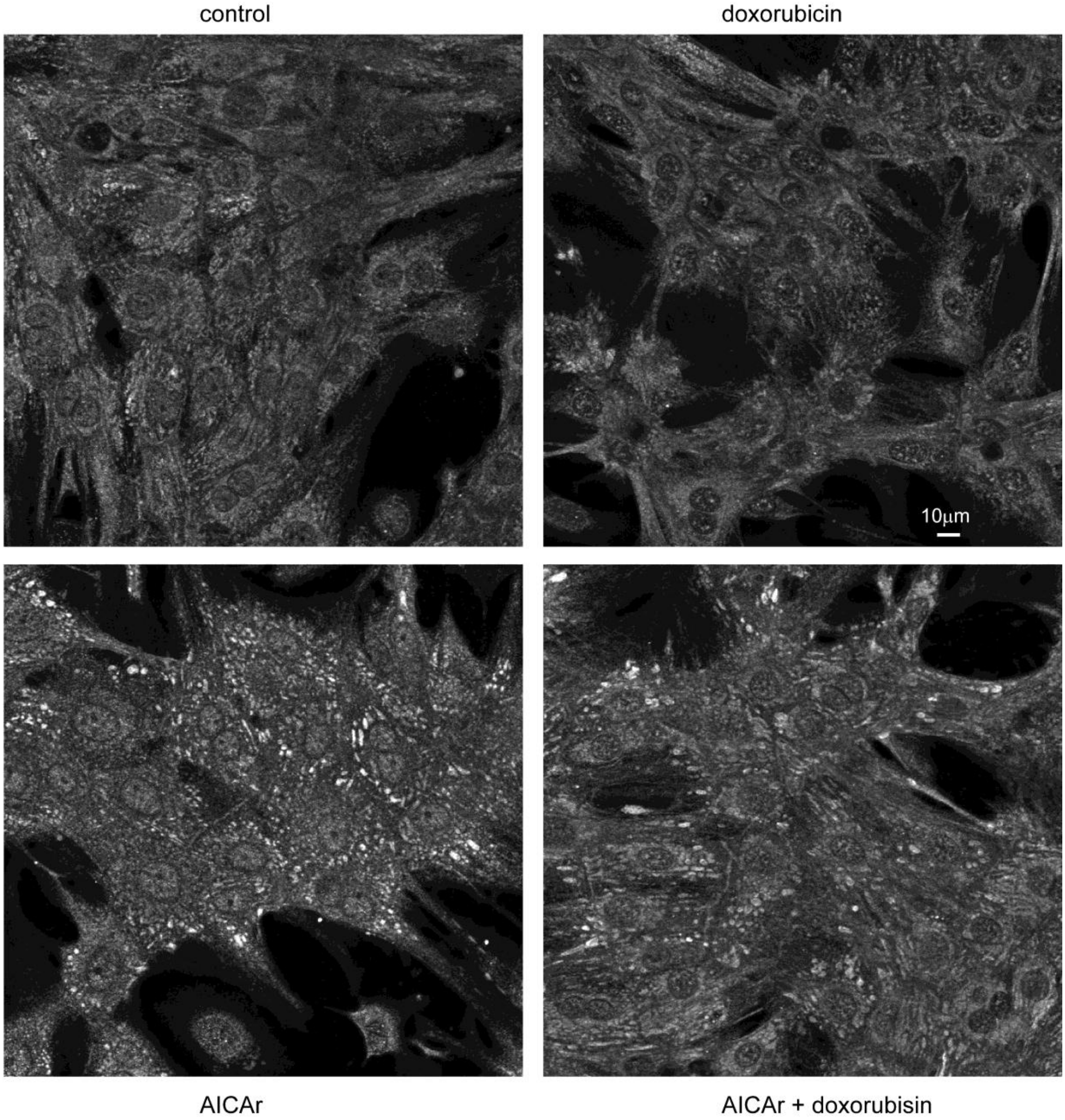


**Supplemental Fig. 5 AICAr and doxorubicin effects on NRF2 nuclear localization**
NRF2 immunofluorescence of NRVMs pretreated for 24 hours with or without AICAr, followed by 6 hours of fresh media with or without doxorubicin.

**Supplemental Fig. 6**

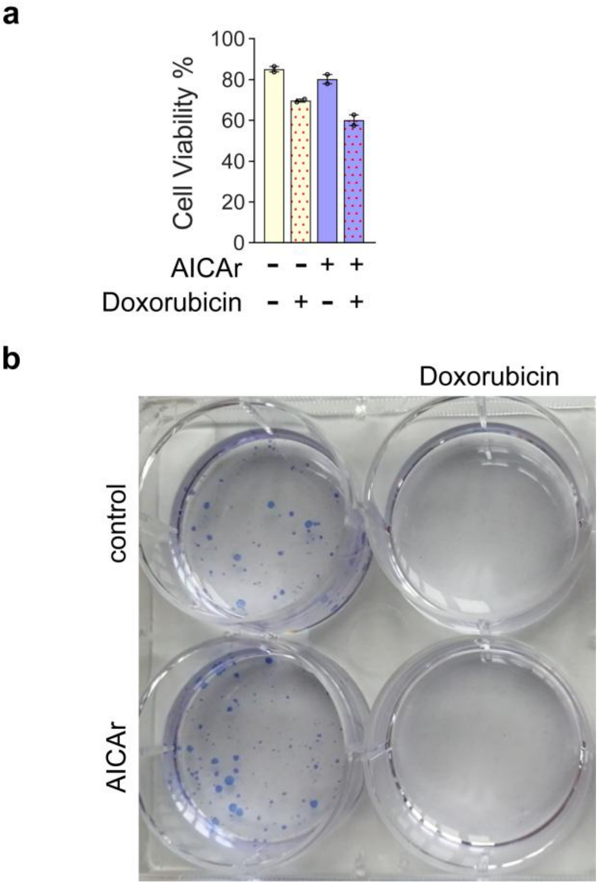


**Supplemental Fig. 6 MCF7 Cells are not protected against Doxorubicin following pre-treatment with AICAr** **a** MCF7 cells were cultured in the presence or absence of 500μM AICAr for 24 hours followed by 2μM doxorubicin for 6 hours. 24 hours later, Cell viability % (Live cells per ml/Total cells per ml) was analyzed via Trypan Blue Assay & CellDrop. Results are expressed as mean +/- SEM. * Indicates $p<0.05$ relative to control. N = 2 independent experiments. **b** The media containing floating cells was removed prior to the Trypan Blue Assay and replated onto a new 6-well plate. The plate was maintained for 10 additional days after which new cell colonies were identified by Coomassie staining
